# Enhanced resistance to bacterial and oomycete pathogens by short tandem target mimic RNAs in tomato

**DOI:** 10.1101/396564

**Authors:** Alex Canto-Pastor, Bruno AMC. Santos, Adrian A. Valli, William Summers, Sebastian Schornack, David C. Baulcombe

**Affiliations:** Department of Plant Sciences, University of Cambridge, Cambridge CB2 3EA, United Kingdom; Sainsbury Laboratory, University of Cambridge, Cambridge CB2 1LR, United Kingdom

## Abstract

Nucleotide binding site leucine-rich repeat (NLR) proteins of the plant innate immune system are negatively regulated by the miR482/2118 family microRNAs (miRNAs) that are in a distinct 22nt class of miRNAs with a double mode of action. First they cleave the target RNA, as with the canonical 21nt miRNAs, and second they trigger secondary siRNA production using the target RNA as a template. Here we address the extent to which the miR482/2118 family affects expression of NLR mRNAs and disease resistance. First we show that structural differences of miR482/2118 family members in tomato (*Solanum lycopersicum*) are functionally significant. The predicted target of the miR482 subfamily is conserved motif in multiple NLR mRNAs whereas, for miR2118b, it is a novel non-coding RNA target formed by rearrangement of several different NLR genes. From RNA sequencing and degradome data in lines expressing short tandem target mimic (STTM) RNAs of miR482/2118 we confirm the different targets of these miRNAs. The effect on NLR mRNA accumulation is slight but, nevertheless, the tomato STTM lines display enhanced resistance to infection with the oomycete and bacterial pathogens. These data implicate an RNA cascade of miRNAs and secondary siRNAs in the regulation of NLR RNAs and show that the encoded NLR proteins have a role in quantitative disease resistance in addition to dominant gene resistance that has been well characterized elsewhere. We also illustrate the use of STTM RNA in a biotechnological approach for enhancing quantitative disease resistance in highly bred cultivars.

## Introduction

Nucleotide binding site leucine-rich repeat (NLR) proteins of plants are central components of the innate immune system that protects against pests and pathogens. These proteins are encoded in multigene families and they regulate signal transduction pathways leading to disease resistance (1). There are several negative regulators of these resistance pathways that, presumably, reduce the likelihood that disease resistance is activated in the absence of a pest or pathogen. Such pathogen-independent induction could be damaging to the host because the disease resistance mechanisms may be associated with programmed cell death called hypersensitive response (HR), changes to the cell wall, production of active oxygen species and strong activation of *pathogenesis-related* (*PR*) genes that compromise reproductive fitness. Transgenic lines overexpressing NLRs (2–4), and gain-of-function mutations like *suppressor of npr1-1, constitutive 1* (*snc1*) (5) or *suppressor of salicylic acid insensitive 4* (*ssi4*) (6) illustrate how inappropriate activation of disease resistance can damage the plant. In normal conditions the negative regulators may mitigate the potential cost of NLR gene diversification and they could be particularly beneficial to plants with many NLR genes (7).

The importance of these negative regulators is reflected in their diversity. There are, for example, suppressor proteins affecting either the NLR proteins themselves or components of the downstream signal transduction pathways (8, 9). There are also micro(mi)RNAs and small interfering (si)RNA negative regulators of NLR mRNA (10–13). These small (s)RNAs bind to their target mRNA by Watson-Crick base pairing and they silence the expression of the mRNA-encoded protein through various mechanisms affecting RNA stability or translation in which the effectors are proteins of the Argonaute family (14).

Amongst the miRNA regulators of NLRs (7) the miR482/2118 family is the most diverse. This family is present in modern seed plants although, in some instances, the apparent conservation may reflect repeated rounds of convergent evolution (15). The miR482/2118 family members are all 22nt rather than the more usual 21nt in length (10–12) and the additional nucleotide is functionally significant because it influences the fate of the targeted mRNA: targets of 21nt miRNA are simply degraded by the Argonaute nuclease activity (14) whereas 22nt miRNA targets are converted into a double stranded RNA by the RNA dependent RNA polymerase 6 (RDR6). The dsRNA is then cleaved by a DCL protein to generate an array of 21nt secondary siRNAs that may be phased with respect to the binding site of the miRNA (16, 17). As a result of this process there is the potential for 22nt miRNAs to establish regulatory cascades in which mRNAs are targeted by both primary miRNAs and secondary sRNAs.

Here we focus on the miR482/2118 family in tomato and we set out to establish the extent to which they influence NLR RNAs and disease resistance. Our approach involved the use short tandem target mimic (STTM) RNAs that would inactivate the miR482/2118 family members and prevent primary or secondary regulation of NLR mRNAs. Our findings confirm that, although the miR482 and miR2118 subfamilies have similar sequences, they are functionally distinct in tomato. The miR482 subfamily targets NLR mRNAs, as already described (10–12) and triggers secondary siRNA production. It is possible, therefore, that NLR RNAs could be both primary and secondary targets of the miR482 family. Only one isoform of miR2118 - miR2118b - is likely to influence NLRs mostly through secondary siRNAs. Its main primary target is not an NLR but a long non-coding RNA – *TAS5* – from which secondary sRNAs are produced that could target NLR coding sequence RNAs. This primary and secondary negative regulation of NLR RNAs by miR482/2118b has an effect on quantitative disease resistance because the STTM lines are less susceptible to an oomycete and to a bacterial pathogen than control lines. This enhanced resistance was achieved without a large effect on growth and development of the plants and it may be useful to protect tomato and other species against pests and diseases.

## Results

**Revisiting the miR482/2118 family and their targets in tomato.** There are numerous families of miRNAs that negatively regulate defence by targeting conserved motifs in NLR RNAs. The miR482/2118 family is the most extensive of these families and it is present in most lineages of seed plants due to either conservation or convergent evolution (7). The present analysis focused on the five members of this family (Table S1) that: (i) align to the tomato genome and (ii) feature in our sRNA datasets from leaves of one-month-old tomato plants.

The sequences of these five miRNAs (Fig. 1A) are complementary to the RNA representation of the conserved P-loop motif (GMGGVGKT) in NLR proteins with sequence variation at 5/6 variable sites corresponding to wobble positions. This pattern indicates that this small family of miRNAs could target a larger number of NLR RNAs with synonymous coding sequence variation (12). Using refined prediction algorithms (18) we estimate that individual miR482/2118 species could potentially target between 15 and 55 NLR RNAs (Fig. 1B; Table S2).

**Fig. 1.**
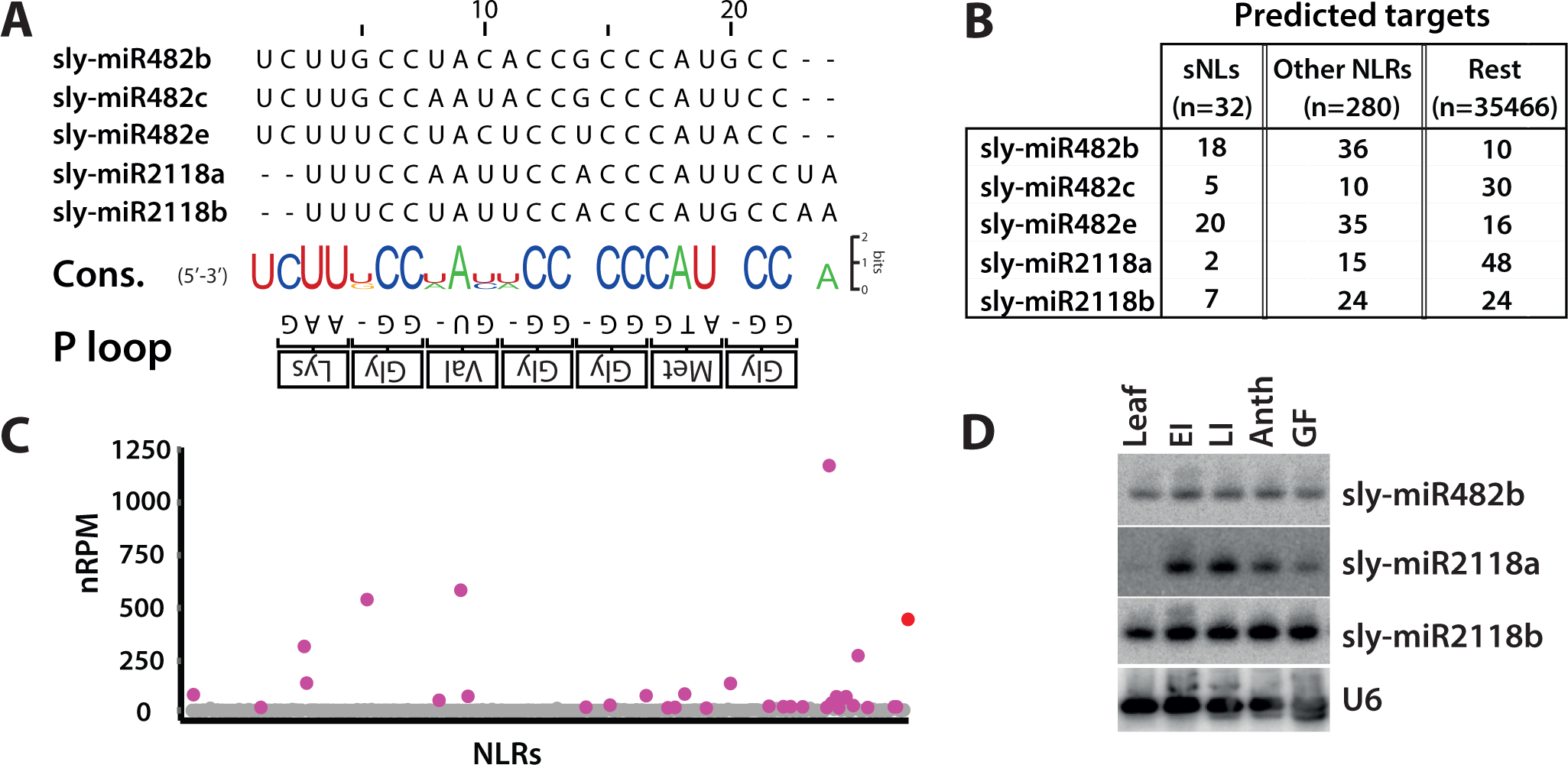
The miR482/2118 family in tomato. (A) Nucleotide sequence alignment of mature miR482/2118 members in tomato. The consensus sequence of each position in the alignment versus the P-loop motif is shown at the bottom. (B) Dot-plot representing the sum of 21-nt sRNA normalized counts per million (nRPM) aligning to an individual NLR. A total of 32 NLRs presented > 10 nRPM counts and were defined as sNLs (purple). *TAS5* (red) is added as a reference. (C) Summary of target prediction of all miR482/2118 members. (D) RNA gel blot analysis of tomato miR482/2118 members in various tissues of plant development. The lower image shows the same blot hybridized with U6, as a loading control. EI: early influorescence; LI: Late influorescence; Anth: Anthesis; GF: Green fruit.

The miR482/2118 family members also have the potential to trigger secondary siRNA production on their mRNA targets (16) because they are 22-nt in length (10–12) and, correspondingly, there were 32 NLR genomic regions with 10 or more normalized 21-22nt siRNA reads per million (nRPM) mapping (Fig. 1C) from our sRNA datasets. Other studies in different species refer to these NLRs with overlapping siR-NAs as phasi-NLRs or pNLs (10, 15, 19) although the extent of phasing may be low. We use the term sNL here referring to NLR mRNAs with secondary sRNAs that may or may not be phased. Most of these tomato sNLs in our datasets are predicted targets of miR482s, with 30 of 32 of them encoding NLRs with coiled-coil (CNLs) rather than TIR (TNLs) domains at the amino terminus (Fig. 1B; Table S3).

Compared to miR482 predictions, potential targets of miR2118s were less enriched for NLRs (Fig. 1B; Table S2) or secondary siRNAs. One of the two isoforms - miR2118a - like its close homologue in other species (20, 21), was much less abundant in leaf tissues than in flower and fruit (Fig. 1D) and it may not have a large effect on NLR mRNAs. In contrast miR2118b, like miR482s, was abundant in leaves and reproductive tissues (Fig. 1D) (12) and it can potentially target several RNAs including 7 NLRs (Fig. 1B; Table S2) and an RNA species that had been identified previously as *TAS5* (22).

Our analysis reveals that *TAS5* is atypical of NLR RNAs because it has sequence similarity to both CNL and TNL types of NLR RNA on both the sense and antisense orientations (Fig. S1). It is unlikely to be translated into a functional protein (calculated coding potential score of -1.146 (23, 24)) and a more likely interpretation is that *TAS5* is a long noncoding RNA comprised of rearranged and degenerated sequences from multiple NLR genes (Fig. S1). It has two miR2118b target sites that were detected bioinformatically and by degradome analysis (Fig. S2, Tables S2&S4) and the siRNAs are predominantly in a phased register. This phasing pattern starts at or adjacent to the more 5’ miR2118b target sites (Fig. S2).

These findings indicate that there are both structural and functional differences between the miR482 and miR2118 subfamilies. The miR482 subfamily (12) targets the CNL NLRs directly and may also have an indirect effect on NLR RNAs via secondary siRNAs. The miR2118 subfamily, in contrast, is different in that its alignment with the P-loop coding motif is shifted by two nucleotides and is only associated with secondary siRNAs (Fig. 1C) in one instance. The miR2118a variant is likely to have little if any effect on NLRs. The miR2118b, in contrast may act via TAS5 secondary siRNAs on both TNL and CNL type NLRs.

**Target mimics block production of secondary NLR siRNAs.** To further investigate the function of miRNAs from the miR482/2118 family in tomato, we generated short tandem target mimic (STTM) transgenes. They were expressed from the strong 35S promoter and were designed so that their transcripts would inactivate the miRNAs 482 or 2118b. We did not test miR2118a as it is expressed predominantly in reproductive tissue (Fig. 1D). The design of the STTM constructs was based on the *IPS1* natural target mimic of miR399 (25) with two tandem target sites (Fig. 2A) to enhance efficiency (26). These target mimic sequences had a three-nucleotide bulge in the miRNA/target RNA duplex structure that would prevent cleavage of the target mimic RNA and channel the miRNAs away from their natural targets.

**Fig. 2.**
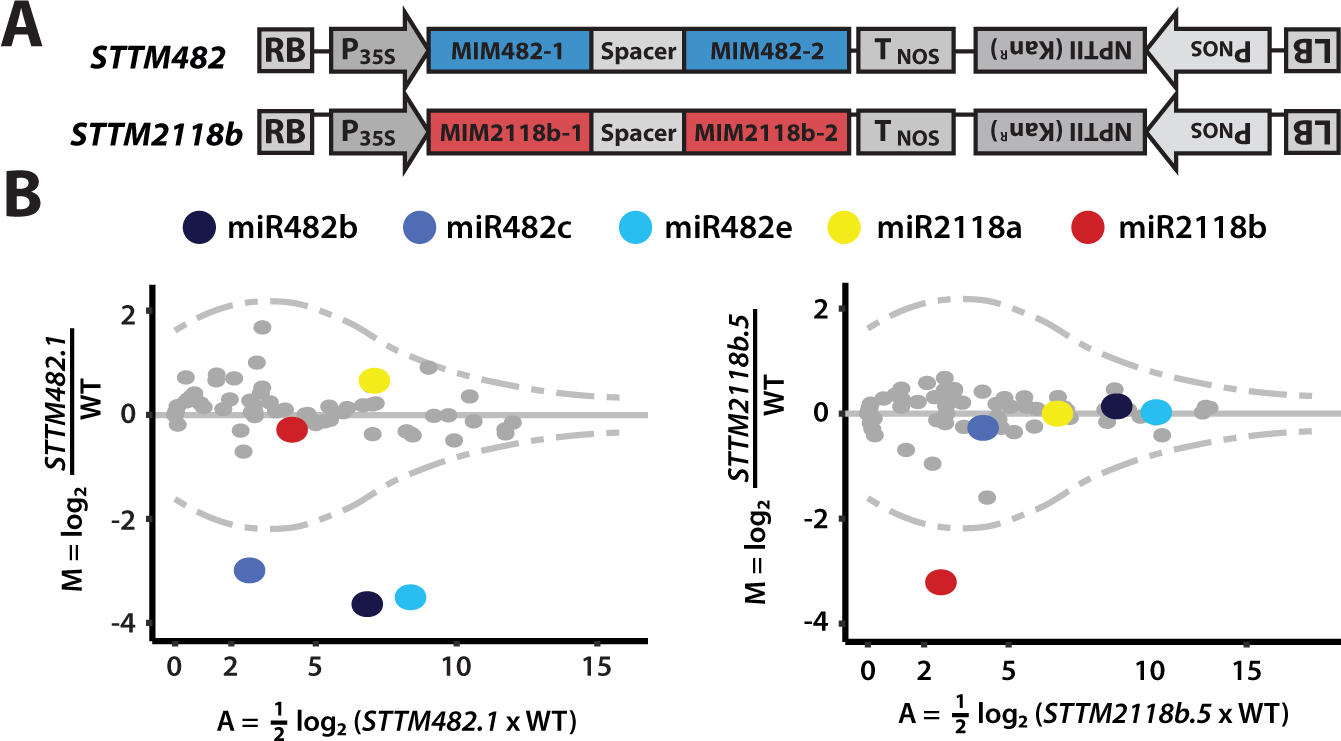
STTMs inactivate specific miRNAs and inhibit production of secondary siRNAs. (A) Diagram of target mimic constructs used in this study. (B) MA plot showing fold changes of miRNAs in STTM lines. Tomato mature miRNA sequences were extracted from miRBASE. The blue dots indicate miR482, yellow is miR2118a and red is miR2118b; grey indicate other miRNAs. The dotted line represents a Poisson distribution with 1 % significance values at the top and bottom of the range, applying the 0 correction (if nreads=0;+1). sRNA reads are normalized to the whole library with reads per million (nRPM) and presented as the mean from three biological replicates.

Lines transformed with the STTM482 and STTM2118b constructs were screened for transgene expression and effects on target miRNAs. Out of 16 stable lines (8 per construct), half showed reduced levels of the corresponding miRNAs (Fig. S3). From these lines, for subsequent detailed analysis, the top inactivating lines for STTM482 (lines 1 & 3) and STTM2118b (lines 5 & 7) were carried to the further generations. In one month-old T2 plants, high throughput sequence analysis of miRNAs in *STTM482.1* revealed a specific tenfold reduction over wild type in all miR482 subfamily members whereas in *STTM2118b.5* the effect was exclusively on miR2118b (Fig. 2B). All other miRNAs were similarly abundant in the STTM lines and wild type plants. The effect of the STTMs, therefore, was specific for the cognate miRNA.

There was an additional effect of the STTMs on sRNAs other than miRNAs. To detect this effect we aligned sRNA reads to the tomato genome and identified differential sRNA loci (DSLs) using the segmentSeq and baySeq packages (27). DSLs in *STTM482.1* were enriched for sNLs and most of them (10/14) overlapped with predicted targets of miR482 members, whereas in *STTM2118b.5* the only DSL was *TAS5* (Table S6). The siRNA levels at 14 DSL sNLs were at least twofold higher in wild type than in *STTM482.1* (Table S7). At *LRR1* (*Solyc02g036270*) and *LRR2* (*Solyc04g005540*), two well-studied CNL RNAs targeted by miR482 (12), there were abundant siRNA aligned to the 3’ side of the miRNA cleavage sites (Fig. 3B) that were four-fold reduced in the *STTM482.1* but not in *STTM2118b.5*. Conversely there was a 5-fold reduction in TAS5-derived siRNAs in *STTM2118b.5* but not in *STTM482.1* (Fig. 3B; Table S7).

**Fig. 3.**
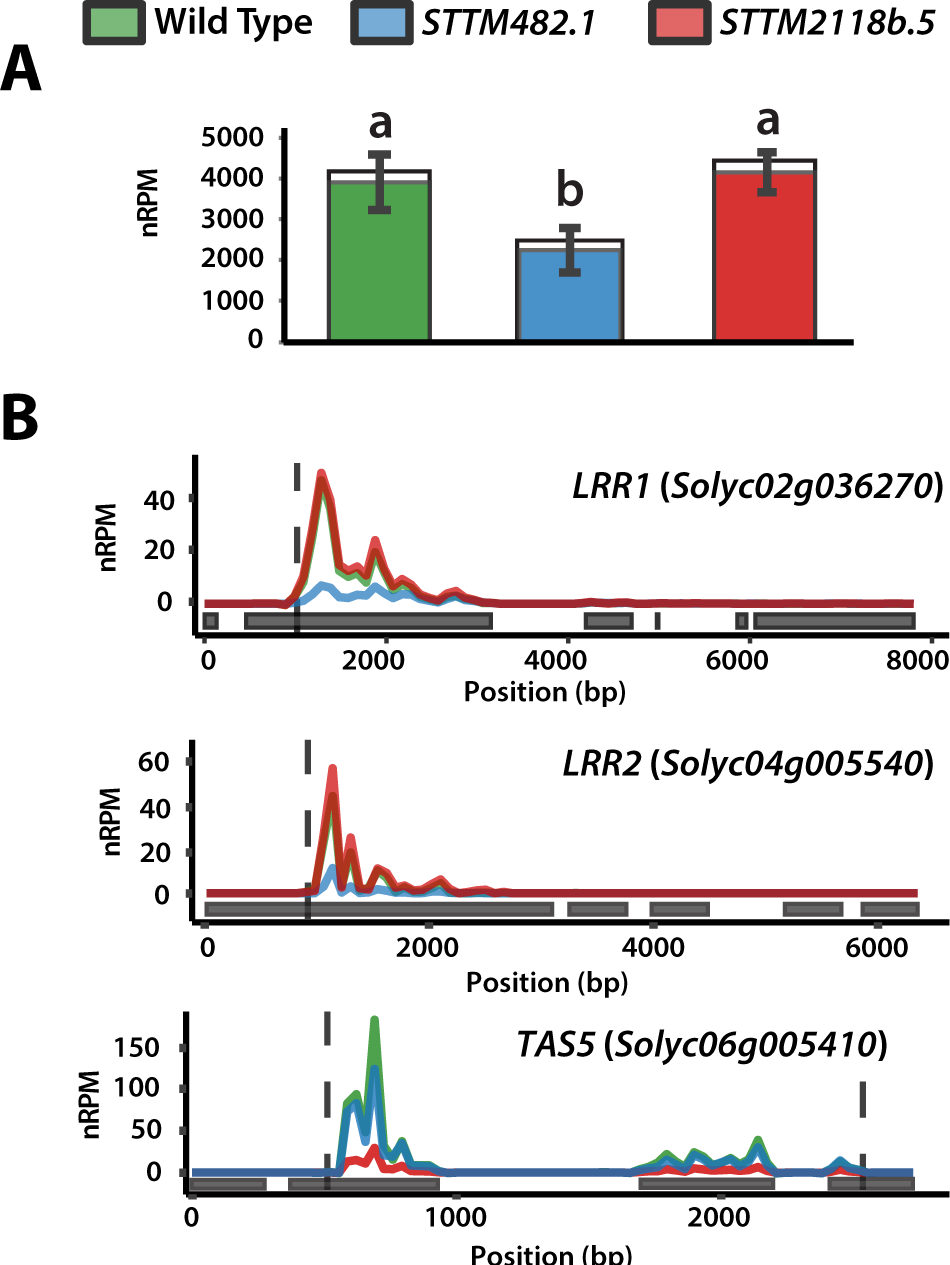
miR482- and miR2118b-mediated silencing is relieved in STTM482 and STTM2118b lines, respectively. (A) Bar-plot of total 21-nt siRNA counts in NLRs in wild type and STTM lines. Coloured fractions correspond to sNLs. Statistically significant differences were found using one-way ANOVA test followed by Tukey HSD at 95% confidence limits. (B) Abundance of 21-nt siRNAs along two sNLs (*LRR1* and *LRR2*, two well-studied CNL genes targeted by miR482) and *TAS5* loci. Positions corresponding to the cleavage sites of their respective miRNA triggers (miR482s and miR2118b, respectively) are indicated by dotted lines. Grey boxes indicate the position of exons. sRNA reads are normalized to the whole library with reads per million (nRPM) and presented as the mean from three biological replicates.

**Target mimic effects on NLR mRNA accumulation.** We further investigated the effect of miR482/2118 species on the NLR mRNA s using RNAseq data of WT and T2 STTM lines. We used young and old leaves and detached leaves inoculated with zoospore droplets of *Phytophthora infestans* 88069. The analysis of these datasets revealed that the overall effects of the STTM RNAs were small in all of the conditions tested. The expressed sNLs were more affected than other NLRs or other genes but, only in the infected plants, was the trend towards elevated expression (Fig. 4A). The previously described miR482 targets *LRR1* and *LRR2* and a few other sNLs and NLR RNAs showed consistent small upregulation (1.5 to 2.5 fold changes) in all conditions in STTM482 but not STTM2118b lines and *TAS5* was consistently upregulated in STTM2118b (Table S7) but not in the STTM482 lines. These observations were validated using qRT-PCR (Fig. S5). To find out whether the small upregulation effect was due directly to reduced targeting by the respective miRNAs we focused on the *P. infestans*-inoculated samples and we compared the targeting score of either miR482 or miR2118b with the degree of overexpression for each of the sNLs and expressed NLRs (Table S2). There was no correlation (Fig. 4B) and we conclude from these data (Fig. 3&4) that the miR482/2118 species are regulators of NLR expression but, to account for the lack of a simple correlation of targeting score and mRNA accumulation, there may be additional mechanisms involved, as discussed below.

**Fig. 4.**
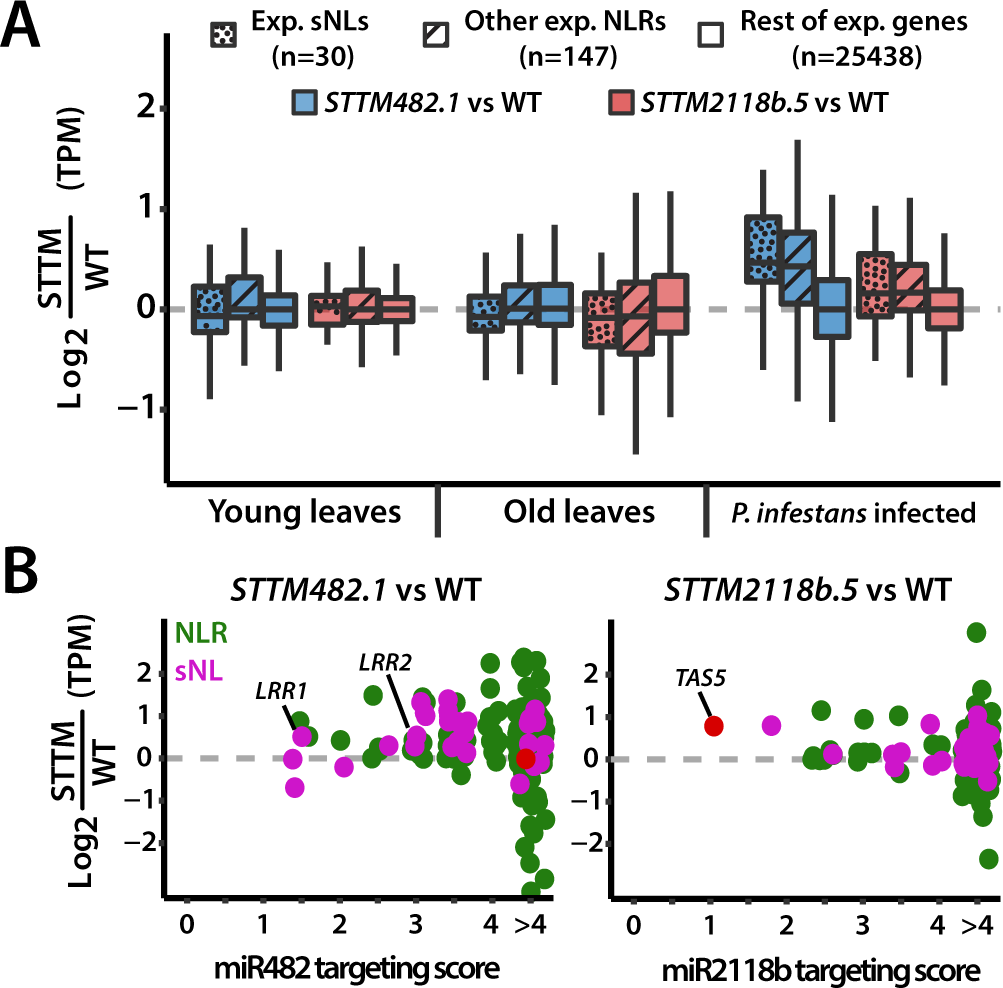
Overall NLR expression is increased in STTM482 and STTM2118b lines during biotic stress. (A) Box-plot of transcript abundances differences between STTM lines and wild type across different conditions. Colours correspond to *STTM482.1* (blue) and *STTM2118b.5* (red) vs wild type. Dotted patterns represent expressed sNLs, while other NLRs are represented as striped boxes, and plain boxes represent the rest of genes in the tomato genome. (B) Scatter plot representing changes in abundance against the targeting score by either miR482 or miR2118b for each of the expressed sNL (violet) or other NLR (green) in STTM lines vs wild type during P. infestans infection. TAS5 (red) was added as a reference. In all instances RNA abundances are calculated as transcripts per million (TPM) and presented as the mean of six biological replicates.

**Inactivation of miR482 and miR2118b enhances resistance to *Phytophthora infestans* and *Pseudomonas syingae***. To investigate the effect of miR482/2118 on disease resistance, we inoculated detached leaves of T2 STTM lines and wild type plants with zoospore droplets of the P. infestans 88069. The tomato cultivar M82 (LA3475), from which all the plants in this study are derived, is highly susceptible to P. infestans (28). We measured the size of necrotic lesions to measure progression of the disease. According to this criterion the *STTM482.1* and *STTM2118b.5* were less susceptible than the non-transgenic control plants with significantly smaller lesion size at 3 days post infection (dpi) (Fig. 5A&B). As a transgene control, a transgenic line expressing a STTM against miR171 (line *STTM171.1*), a miRNA with no reported role in plant defence, was used and was as susceptible as non-transgenic lines.

**Fig. 5.**
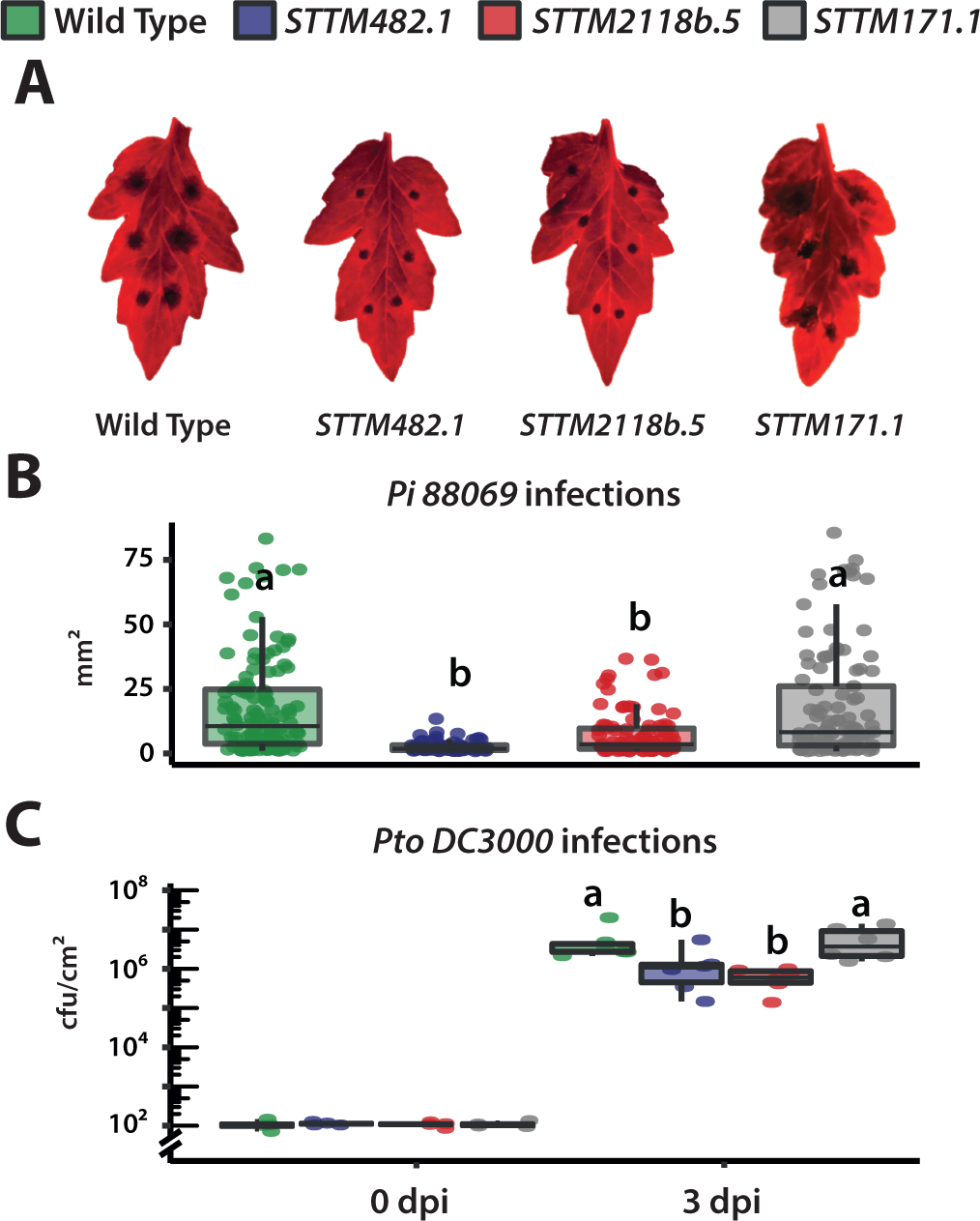
Sequestration of miR482 and miR2118b influences broad-spectrum resistance to pathogens. (A) Representative images of detached leaves under blue light 3 days post infection with *Phytophthora infestans Pi 88069*. (B) Boxplot and leaf images of lesion size in WT and mimicry lines. Statistically significant differences were determined using one-way ANOVA test followed by Tukey HSD at 95% confidence limits (n=12). (C) Boxplot of bacterial population in WT and STTM lines leaves infected with *Pseudomonas syringae pv. tomato DC3000*. Bacterial counts at 0 and 3 days post leaf infiltration. Statistically significant differences were determined using ANOVA test followed by Tukey HSD at 95% confidence limits (n=6). Both infection experiments were repeated three times with similar results.

We also tested the effect of the STTM constructs on resistance against *Pseudomonas syringae pv. tomato DC3000*. Bacterial titers in inoculated leaves attached to the plant were lower at 3 days post inoculation (dpi) in both *STTM482.1* and *STTM2118b.5* than wild type and *STTM171.1* lines (Fig. 5C). This effect indicates that the primary or secondary targets of miR482s and miR2118b have a role in anti-bacterial immunity.

Defense activation may be at the expense of the plant’s fitness (29). In the STTM lines, however, general growth was similar in the control and STTM lines used in the disease resistance tests other than a small decrease in shoot length in *STTM482.1* (Fig. S6). These tests may not be sensitive to other slight differences in fitness but they indicate that enhanced disease resistance in the STTM lines was not associated with gross effects on growth and development.

**miR482/2118b as regulators of innate immunity.** In this paper we describe definitive evidence that miR482/2118b family members are negative regulators of innate immunity in tomato. The miR482 subfamily members were known previously to affect the level of NLR RNAs and their overexpression resulted in hyper-susceptibility to *Verticillium* and *P. infestans* consistent with a role in immunity (30, 31). Now, by suppression of miR482 and miR2118b using STTMs, we enhance resistance against both *P. infestans* and *Ps. syringae* and provide direct evidence that the miR482/2118b family is a negative regulator of innate immunity (Fig. 5). Based on the target site specificity of miR482s and miR2118b it is highly likely that resistance is due to effects on NLRs. Our findings confirm the very recent report that STTM482b confers enhanced resistance against *P. infestans* (31).

Well-described mechanisms involving siRNA and miRNA result in cleavage and degradation of the target RNA (14) and we had expected that the sNL RNAs and other NLR RNA targets of miR482/2118b family would increase in abundance in the STTM lines. There was, however, only a small increase (Table S7; Fig. 4&S6) and, to reconcile these RNAseq data with the enhanced resistance, we hypothesize that there may also have been a translational effect that would not have affected the abundance of the encoded NLRs. To further enhance the resistance it may be necessary to use miRNA knockout rather than STTM transgenes that do not give complete suppression of the miRNAs and their effects (Fig. 2B&3).

NLR genes are normally associated with dominant gene resistance and mechanisms that confine the pathogen to the site of initial infection. This dominant gene resistance was the first described phenotype of NLRs (32) and it is normally specific for races or strains of the pathogen. The NLRs in this race-specific resistance mediate direct or indirect recognition of effectors that are transported into the infected cell (1). This “effector-triggered immunity” then triggers metabolic and molecular changes that hinder the progression of disease (33). Our tests were not, however, with race-specific resistance. The plant genotypes were fully susceptible to both pathogens and the STTM-mediated effect was manifested as quantitative resistance rather than complete suppression of the pathogen (Fig. 5).

To explain this quantitative resistance in the STTM plants, we propose that there is a low level recognition of pathogen effectors even in susceptible plants but that any triggering of resistance would be too slow or too weak to prevent accumulation and spread of *P. infestans* and *Ps. syringae*. In the STTM plants, however, the block on silencing of NLRs may allow stronger recognition and more rapid activation of defense. Alternatively, any increased expression of NLRs in the STTM lines could cause auto-activation of disease resistance in the absence of elicitor recognition analogously to the effects of overexpressing NLRs in non-infected plants (2–4). How many NLRs affect the quantitative resistance in the STTM plants? With miR482 there are at least 32 NLRs generating secondary siRNAs (Fig. 1) and, at least in principle, there could be many more implicated in a miR482/2118b cascade. The first layer in this cascade would involve NLR RNAs that are direct targets of miR482 and a second layer might involve NLR targets of secondary siRNAs. At each layer there could be posttranscriptional or translational suppression of NLR target RNAs.

With miR2118b, however, the main primary target generating secondary siRNA is a non-coding RNA – *TAS5*. It is unlikely that this *TAS5* RNA affects resistance directly and a feasible interpretation of the enhanced resistance (Fig. 5) involves indirect effects mediated by the secondary siRNAs that are reduced in the *STTM2118b.5* (Figures 3&S2; Tables S4-7). These *TAS5* secondary siRNA targets, as with the primary miR482/2118b targets, could be subject to post-transcriptional or translational regulation.

**Evolutionary dynamics of NLR regulation by miRNA.** The miRNA regulation of NLRs is highly variable in angiosperm species. In addition to the miR482/2118 family there are numerous other miRNAs including miR5300, miR6024, miR6026, miR825 and others with the potential to target NLRs (34). All of these miRNAs are variably present, depending on the species, indicating a dynamic evolutionary process in which RNA silencing of NLR RNAs is lost and gained (7, 35).

The miR2118 family is a special case of NLR regulation that is dependent on two genome events specifically in the *Solanaceae* (Fig. S7). One of these changes involves expression of a miR2118b in the vegetative phase including leaves. In most other monocot and dicot species the miR2118, like miR2118a in tomato (Fig. 1) is specific to reproductive phases (20, 21). The second *Solanaceae*-specific genome change (Fig. S1) would have been a rearrangement of TNL and CNL NLR genes resulting in the *TAS5* locus. These changes illustrate how the evolutionary dynamics of NLR regulation are not only dependent on loss or gain of the miRNA genes. There can also be neo-functionalisation of miRNA genes, as with miR2118b, and the addition of noncoding RNAs into the RNA silencing cascade, as with *TAS5*. We have speculated previously that the miR482-mediated down regulation of NLRs is a process that allows the plant to trade-off the costs and benefits of NLRs (35). Accumulation of these NLR gene products has a cost that is presumably related to metabolic changes associated with disease resistance and a benefit due to protection against pests and pathogens. The balance of costs and benefits depends, presumably, on the ecological niche occupied by the plant and on the effectiveness of other defense systems.

In these terms, to explain the apparent minimal cost of STTMs on growth and development of the plants, we suggest that the protected glasshouse environment of our tomato plants allowed them to tolerate the increased expression of NLRs without effects on growth and development of the plant (Fig. S6). There may also be other changes associated with the domestication and breeding of cultivated tomato that protect the plant against the costs of increased NLRs. Further testing will be required to establish whether there is a cryptic growth phenotype associated with STTM expression and also whether the basal immunity can be enhanced further. It could be that a combination of the two STTMs would have a stronger effect than with the individual species. Alternatively the enhanced basal immunity would be useful if it is part of an integrated management strategy to protect crops against disease.

## Materials and methods

**Plant strains and growth conditions.** Tomato (*Solanum lycopersicum*) cultivars M82 were raised from seeds in compost (Levington^™^ M3) and maintained in a growth room with 16/8h light/dark periods at 22°C (day) and 18°C (night), with 60% relative humidity, at a light intensity of 300 µmol photons m^-2^ ·s^-1^. Agrobacterium tumefaciens-mediated stable transformation of tomato plants were performed based on published work (36).

**Cloning and vector construction.** The STTMs vector construction was done based on a previous report (26). In brief, a long (110bp) DNA oligo containing two mimic sequences separated by a spacer was designed and cloned into a pENTR L1L2 vector (Invitrogen). The insert in this plasmid (pENTR-STTM) was then LR recombined into pGWB402 destination vector containing a *2X35S* promoter driving the expression of the insert, and kanamycin-resistant marker (*NOS promoter:NPTII:NOS terminator*) for selection. All constructs were confirmed by Sanger sequencing.

**Small RNA northern blot.** Small RNA detection was performed using the northern blot technique. In brief, 5 µg of total RNA per sample were prepared in 10 µl, added equal volume of 2X loading buffer (95% deionized formamide, 18 mM EDTA, 0.025% SDS, xylene cyanol FF, bromophenol blue), and heated at 65°C for 5 min. Then placed in ice for 1 min and loaded and run in a 15% polyacrylamide 7 M urea gel, using 0.5X TBE running buffer. RNA was then transferred to a positively charged nylon membrane (Amershan Hybond-N+, GE Healthcare^™^) using overnight capillary system: gels were soaked for 10 min in 20X SSC, then placed on a clean glass plate, membrane on top, 2 pieces of 3MM paper soaked in 20X SSC, and finally 3-5cm of thick paper on top. Another glass plate was put on top and 1kg of weight over this plate. RNA was cross-linked two times per side with 0.12 J of UV light in a Stratalinker^®^ (Agilent^™^). Oligonucleotides and Locked Nucleic Acid (LNA^™^; by Exiqon) probes radiolabelled with *γ*^32^ P-ATP were hybridized in ULTRAhyb-Oligo buffer (Thermo Fisher Scientific^™^) for 12 hours at 40°C or 2 hours at 57°C, respectively. Then washed three times with 2X SSC, 0.2% SDS. Phosphoimager plates (Fujifilm) were exposed and then imaged with a Typhoon 8610 (Molecular Dynamics).

**Target prediction.** All tomato miRNA mature sequences were downloaded from miRBase (v21) (37). MicroRNAs targeting genes were predicted by psRNATarget v1 (Dai and Zhao, 2011). Cut-off values were established based on the optimal scores reported in previous studies (18). Tomato NLR sequences were retrieved from a previous study (38) and curated using the ITAG3.2 annotation.

**Degradome (PARE) analysis.** Parallel analysis of RNA ends (PARE) was performed using the software sPARTA (39). The analysis was done on publicly available datasets from the tomato degradome data of leaf samples (40).

**sRNAseq analysis.** Small RNAs libraries were prepared using the NEBNext ^®^ Small RNA Library Prep (New England Biolabs). In brief, three biological replicates each of 1 month old tomato leaf RNA were prepared using 1 µg of total RNA per sample. After preparation, size selection of libraries was performed using BluePippin 3% agarose cassettes (Sage Science). Each library was barcoded, pooled, and sequenced using a single NextSeq 500/550 High Output Kit v2 (75 cycles). Sequences were demultiplex, and trimmed and filtered using Trim Galore! (Babraham Bioinformatics) with default parameters and reads were concordantly aligned to the Heinz genome SL3.00 version using Bowtie v1.2.0 with modifiers -v 1 -m 50 –best –strata. Identification of sRNA loci and differential expression was performed using the segementSeq package and baySeq respectively (27, 41).

***Phytophthora infestans* infections.** The *Phytophthora infestans* strain in this study is 88069 (42). Cultures were stored in liquid nitrogen and grown on rye sucrose medium. Infection assays were performed on detached tomato leaves, measuring lesion sizes. In brief, four well developed leaves per plant and four plants per condition were detached from fourweek-old plants and placed on water-saturated paper in a tray. Spore suspensions of P. infestans were prepared by rinsing two-week-old plates covered with mycelium with cold water and incubating the sporangiophore at 4°C for 1-2 hours. After release of zoospores, the concentration was adjusted to approximately 5 · 10^4^ spores·ml^-1^. *P. infestans* were spot-inoculated on the abaxial side of the leaf, by placing six 10 µl droplets on various locations right and left of the midvein. The trays were covered and incubated at room temperature at constant light a photoperiod. Disease assessments were performed daily from 3 to 7 days post inoculation (dpi) under blue light using a DarkReader ^®^ Transilluminator (Clare Chemical Research) and a Nikon COOLPIX P520. Lesion diameters were measured using ImageJ software, followed by statistical analysis and plotting in R.

***Pseudomonas syringae* infections.** The *Pseudomonas syringae pv. tomato* strain used in this study is DC3000, which is a pathogen of tomato developed in 1986 as rifampicin-resistant derivative of Pst DC52. Cultures were stored in liquid nitrogen and grown on King’s B medium. Infection assays were performed in planta, inoculating mature tomato leaves and measuring bacterial growth based on previous work (43). Statistical analysis and plotting were done in R.

**RNAseq analysis.** RNAseq libraries were prepared using the Truseq^®^ mRNA HT kit (Illumina). In brief, total RNA from three different conditions (young: 3 weeks old leaves, old: 6 weeks old leaves, and infected: 3 day post inoculation of detached leaves with *P. infestans*) with six biological replicates each of 3 week old, and 6 week old tomato leaves, RNA were prepared using 1 µg of total RNA per sample. PolyA bead selection and strand-specific RNA-seq libraries were made and indexed according to manufacturer instructions. Finalized libraries were sequenced as a pool on one lane of a NextSeq 500/550 High Output Kit v2 (75+75 cycles). Sequences were de-multiplex, and trimmed and filtered using Trim Galore! (Babraham Bioinformatics) with default parameters. Trimmed reads were pseudo-aligned to ITAG3.2 transcriptome using Kallisto (44), with the parameter -b 100. Differential expression was performed on kallisto-estimated counts using the Bioconductor package DESeq2 (45). For visual representations and analysis, abundances were reported as quantile-normalized transcripts per million (TPM). Processing, analysis and plotting were done in R.

**Phylogenetic analysis.** BLASTN analyses were performed using genomic sequences of tomato genes and miRNA precursor sequences against the genomes of all plant model organisms and all available genome assemblies of major Solanaceae species. The threshold expectation value was defined at 10^-3^ to filter out any spurious hits. Any hits were then manually curated.

## ACKNOWLEDGEMENTS

The authors would like to thank James Barlow, Mel Steer, Antonia Yarur and Pawel Baster for technical and horticultural assistance; Stuart Fawke for maintaining and providing *P. infestans 88069*; Zhengming Wang for the *STTM171.1* seeds; and Thomas Hardcastle for bioinformatics advice. This work was supported by the Balzan Foundation and the European Research Council Advanced Investigator Grant ERC-2013-AdG 340642. D.C.B. is the Royal Society Edward Penley Abraham Research Professor. S.S is funded by Gatsby Foundation Fellowship GAT3395/GLD and a Royal Society University Research Fellowship UF160413. The funding bodies had no roles in the design of the study, in the collection, analysis, interpretation of data, or in the writing of the manuscript.

## AUTHOR CONTRIBUTIONS

A.C.P., S.S. and D.C.B. designed research; A.C.P. performed research and analysed data; B.AMC.S assisted with bioinformatics analysis; A.A.V and W.S. contributed new analytic tools; A.A.V. assisted with the writing and analysis, and A.C.P., S.S. and D.C.B. wrote the paper.

## COMPETING FINANCIAL INTERESTS

The authors declare that they have no competing interests.

## Supplementary images

**Fig. S1.**
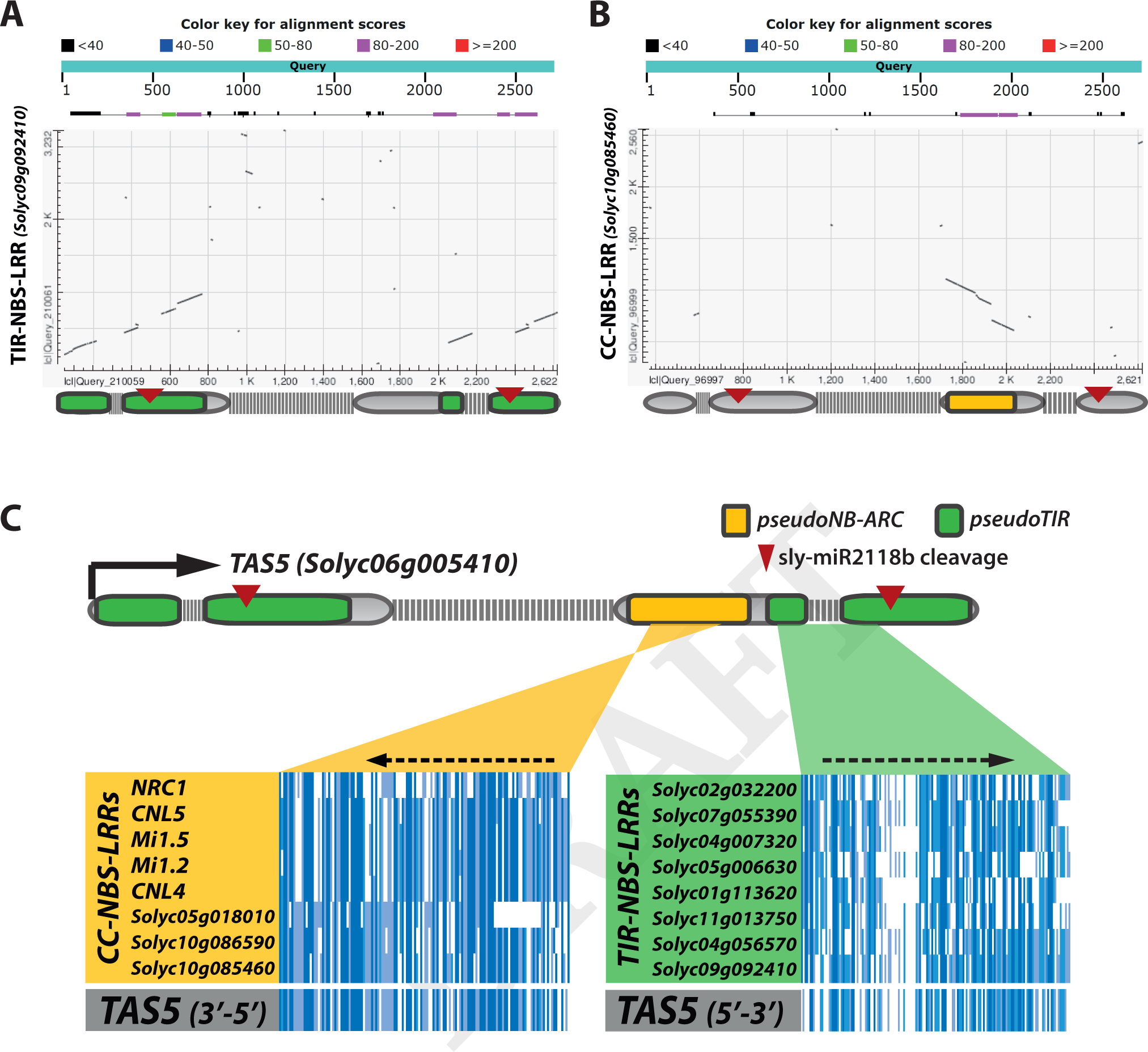
Sequence similarity between NLR domains and TAS5. BLAST (Basic Local Alignment Search Tool) summary and dot plot matrix of TAS5 versus (A) the closest TNL and (B) the closest CNL in the tomato genome. Gene diagram of TAS5 locus is placed underneath the matrix indicates in (A) similarity to a TIR domain in green and (B) similarity to a NB-ARC in yellow. (C) Regions within exons with most significant sequence similarity with known NLR domains are highlighted in yellow and green. Nucleotide sequence alignment of these regions and known tomato NLRs are shown below. The degree of conservation for each nucleotide along the region is represented by the colour, with a dark blue denoting a high level of conservation and a light blue denoting a low level. Dotted arrows indicate direction of the sequence similarity.

**Fig. S2.**
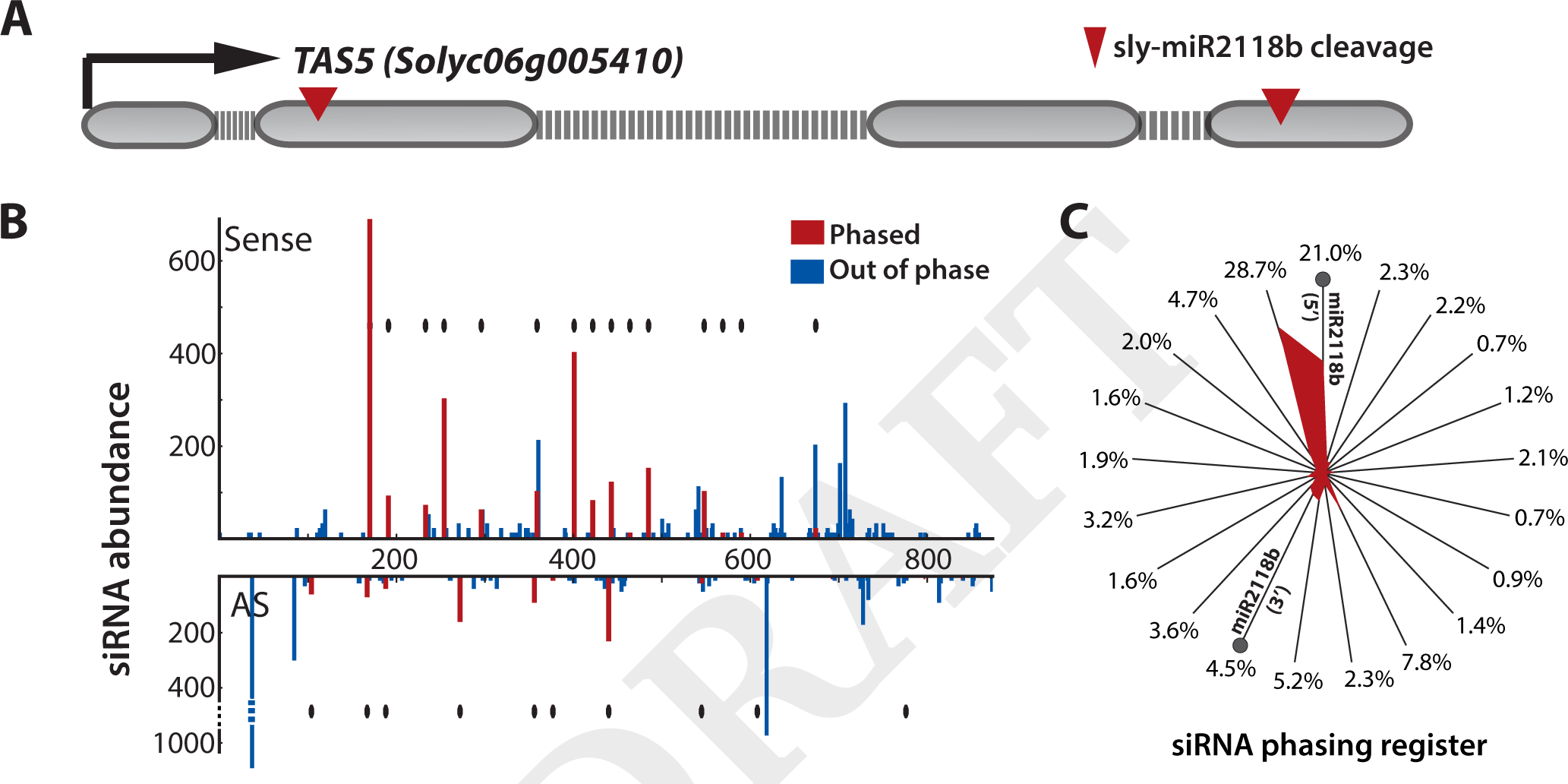
TAS5 is a phasiRNA-producing locus. (A) Gene diagram of TAS5 locus. A 2.7kb region containing 4 exons (grey boxes) and 3 introns (grey dotted lines). The arrow on the left indicates transcription start site and direction. (B) Number of sequenced small RNAs with 5’ residues at each position between the cleavage sites of the TAS5 transcript. Red bars indicate phased sequences while blue indicate out of phased. Red diamonds indicate that expected phased siRNA is present in the sample. (C) Distribution of the phasing of small RNAs at the TAS5 locus. Each spoke of wheel the represents 1 of the 21 possible registers, with the percentage of small RNAs mapping plotted as distance from the centre (correction of 2-nt 3’ overhangs of DCL cleavage was applied when assigning register in the anti-sense strand). The specific registers predicted from 5’ and 3’ cleavage sites of miR2118b are indicated with grey circles.

**Fig. S3.**
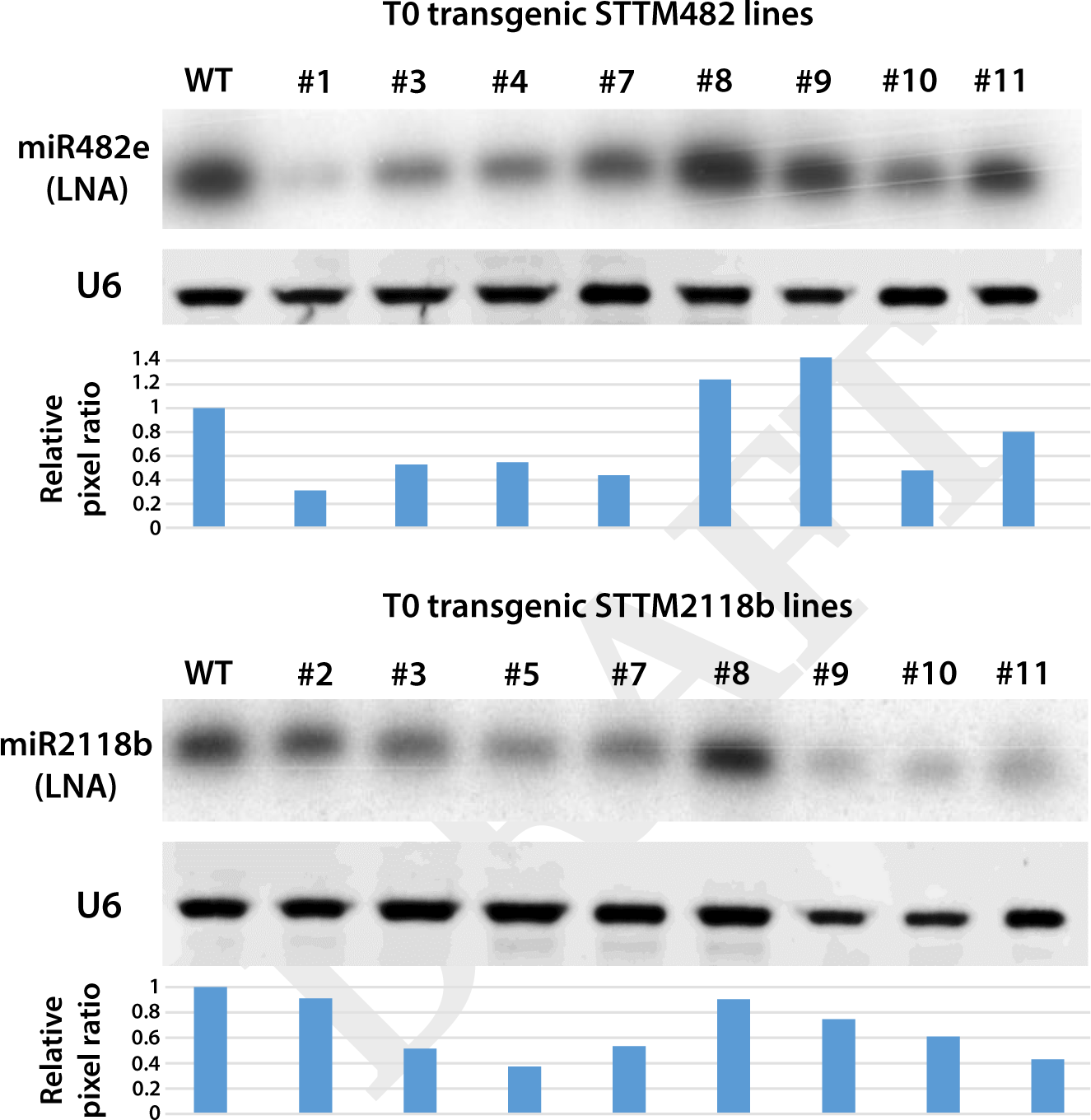
Levels of target miRNA sequestration in first generation STTM transgenic lines. RNA gel blot analysis of tomato transgenic mimic lines. Upper line shows miRNA blotted with highly specific locked nucleic acid (LNA) probes. Lower image shows the same blot hybridized with U6, as a loading control. Barplot indicates relative pixel ratio of miRNA signal vs U6 signal.

**Fig. S4.**
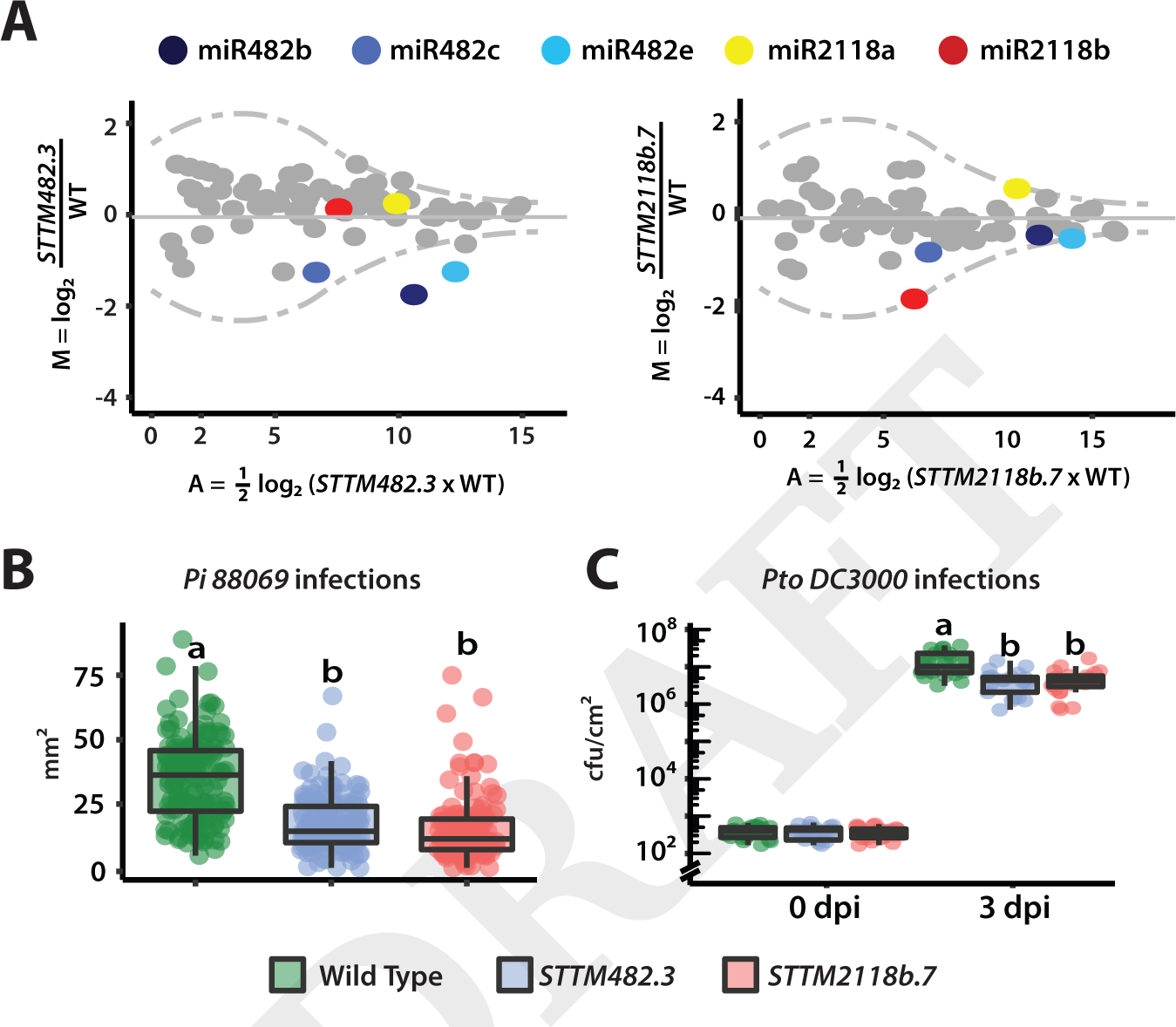
Effects were consistent across independent STTM lines. (A) MA plot showing fold changes of miRNAs in STTM lines. Tomato mature miRNA sequences were extracted from miRBASE. The blue dots indicate miR482, yellow is miR2118a and red is miR2118b; grey indicate other miRNAs. The dotted line represents a Poisson distribution with 1 % significance values at the top and bottom of the range, applying the 0 correction (if nreads=0;+1). sRNA reads are normalized to the whole library with reads per million (nRPM) and presented as the mean from three biological replicates. (B) Boxplot and leaf images of lesion size in WT and mimicry lines. Statistically significant differences were determined using one-way ANOVA test followed by Tukey HSD at 95% confidence limits. (C) Boxplot of bacterial population in WT and STTM lines leaves infected with *Pseudomonas syringae pv. tomato DC3000*. Bacterial counts at 0 and 3 days post leaf infiltration. Statistically significant differences were determined using ANOVA test followed by Tukey HSD at 95% confidence limits.

**Fig. S5.**
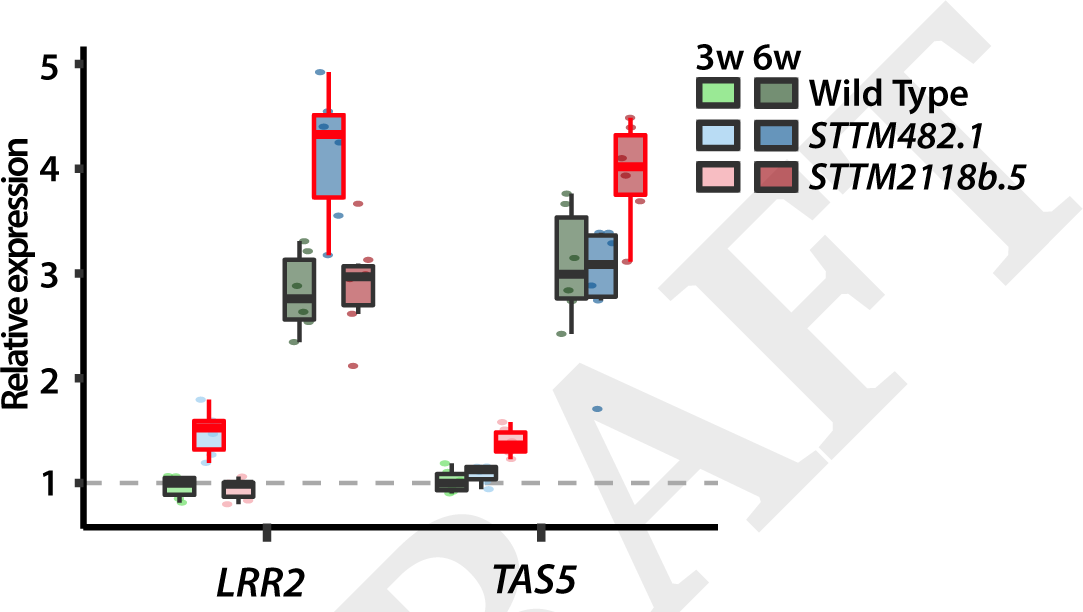
qRT-PCR analysis validates RNA-seq results for miR482 and miR2118b targets. Quantitative PCR analysis for the abundance of target mRNAs LRR2 and TAS5 in 3 and 6 week old leaf tissue (n=6). Expression values were adjusted to tomato housekeeping gene EXP and shown relatively to WT values. Statistically significant differences were explored using two-way ANOVA test followed by Tukey HSD at 95% confidence limits.

**Fig. S6.**
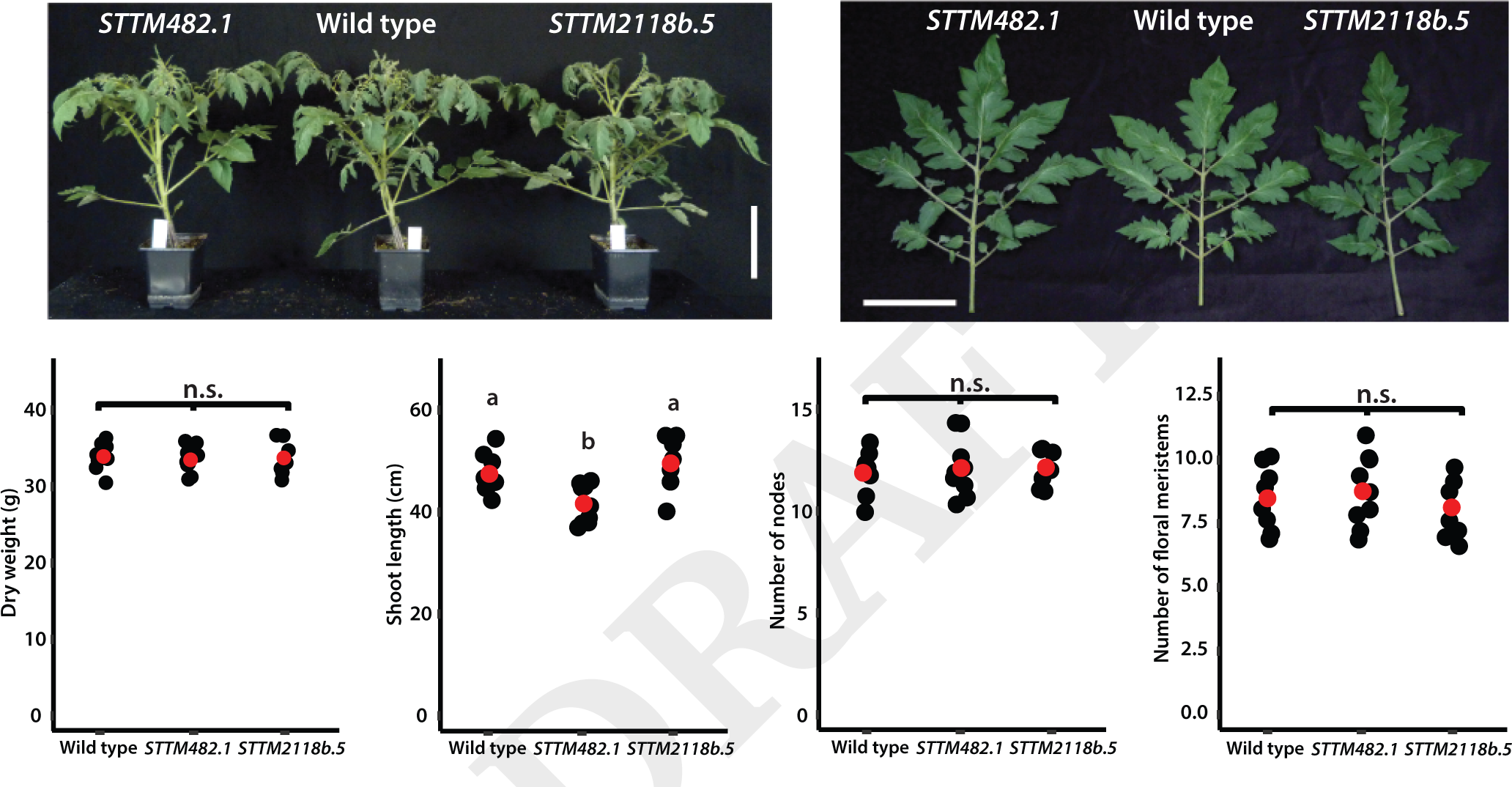
Growth remains majorly unaffected in STTM transgenic lines. (Top) representative images of leaves and whole plants. White scale bars represent 10cm. (Bottom) Dot plots representing differences in shoot dry weight, length, number of nodes, and number of floral meristems. Black dots represent individual plants, and red dots represent means of all biological replicates (n=8). Images and data were collected at 8 week after germination. Statistically significant differences were determined using a two-way ANOVA test followed by Tukey HSD at 95% confidence limits.

**Fig. S7.**
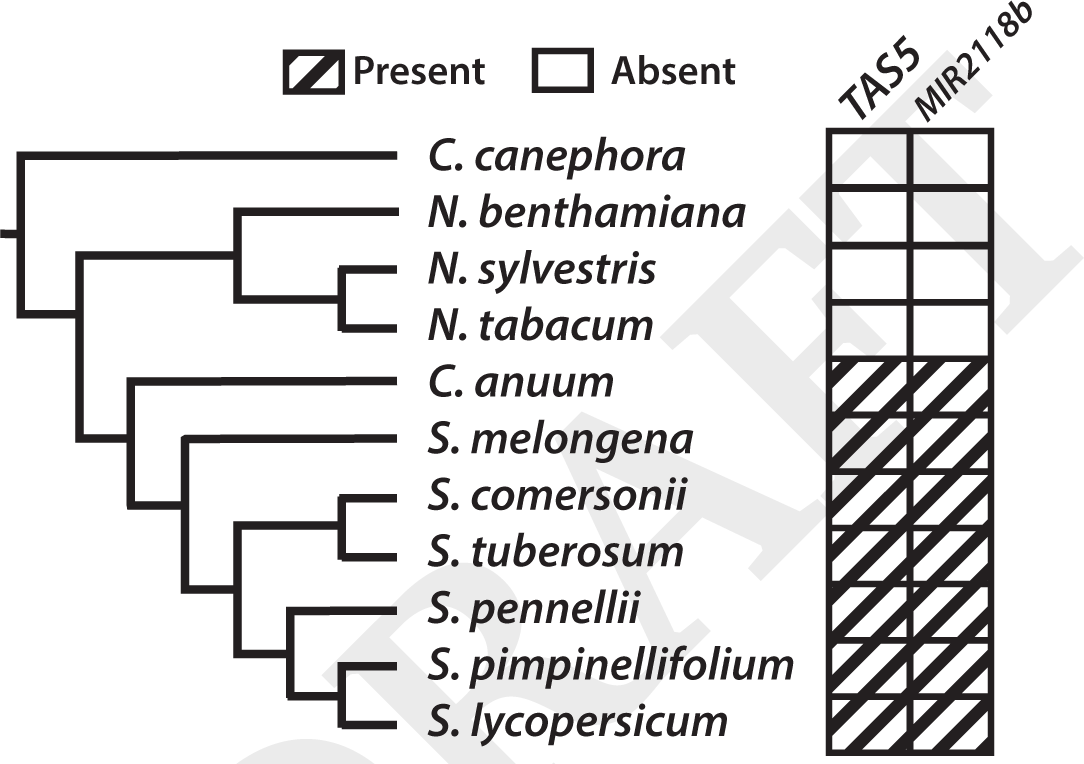
TAS5 and MIR2118b are only present in Solanum species. Diagram summarising the presence or absence of genomic sequences matching TAS5 and MIR2118b in Solanaceae and a close relative. A close sequenced relative of Solanaceae, *Coffea canephora*, was included as an outgroup.

## Supplementary tables

**Table S1.**
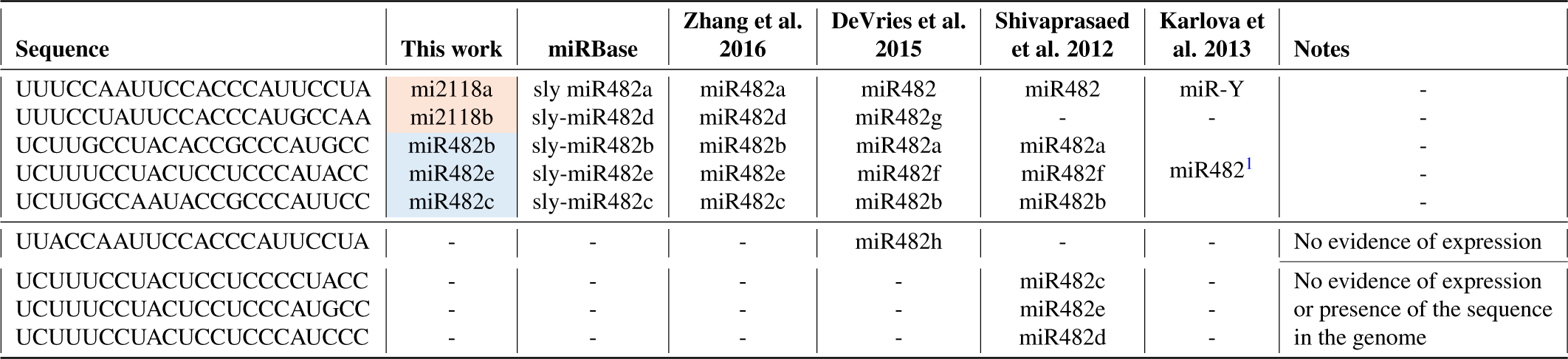
Summary of identified members of the miR482/2118 family in tomato. All available nomenclatures of miR482/2118 members in tomato in the different studies, present in the current literature.

**Table S2.** Summary of all predicted targets of miR482/2118 members.

| miRNA | Target | Score | UPE | Posit | Target description |
| --- | --- | --- | --- | --- | --- |
| mi2118a | Solyc11g008140 | 1 | 11.101 | 1224 | Pectate lyase |
| mi2118a | Solyc01g102920 | 2 | 14.713 | 635 | Disease resistance protein (TIR-NBS-LRR class) |
| mi2118a | Solyc04g007320 | 2 | 21.199 | 1110 | Disease resistance protein (TIR-NBS-LRR class) |
| mi2118a | Solyc01g020371 | 2.5 | 20.058 | 349 | GRF zinc finger family protein |
| mi2118a | Solyc03g116360 | 2.5 | 15.966 | 2405 | Regulator of chromosome condensation (RCC1) family protein |
| mi2118a | Solyc04g024950 | 2.5 | 15.862 | 4 | MATH domain/coiled-coil protein |
| mi2118a | Solyc04g049780 | 2.5 | 13.961 | 70 | Retrovirus-related Pol polyprotein from transposon TNT 1-94 |
| mi2118a | Solyc04g053070 | 2.5 | 17.989 | 278 | DNA topoisomerase |
| mi2118a | Solyc06g009533 | 2.5 | 18.082 | 77 | Kinase family protein |
| mi2118a | Solyc06g062440 | 2.5 | 22.729 | 593 | Disease resistance protein |
| mi2118a | Solyc10g007065 | 2.5 | 17.785 | 580 | Phenylalanyl-tRNA synthetase alpha chain |
| mi2118a | Solyc01g066020 | 3 | 25.484 | 656 | disease resistance protein (TIR-NBS-LRR class) |
| mi2118a | Solyc01g087200 | 3 | 19.95 | 515 | Disease resistance protein |
| mi2118a | Solyc01g090860 | 3 | 8.404 | 782 | Nucleotidyltransferase family protein |
| mi2118a | Solyc02g030100 | 3 | 17.13 | 3745 | Vacuolar protein sorting-associated protein 54 |
| mi2118a | Solyc02g030105 | 3 | 17.13 | 2139 | Vacuolar protein sorting-associated protein 54 |
| mi2118a | Solyc02g091890 | 3 | 6.22 | 4704 | myb-like protein X |
| mi2118a | Solyc03g083130 | 3 | 16.198 | 1745 | gamma-irradiation and mitomycin c induced 1 |
| mi2118a | Solyc05g009750 | 3 | 17.541 | 164 | NBS-LRR resistance protein-like protein |
| mi2118a | Solyc08g005510 | 3 | 18.753 | 1512 | disease resistance protein (TIR-NBS-LRR class) |
| mi2118a | Solyc09g090390 | 3 | 18.271 | 1709 | 2-oxoglutarate-dependent dioxygenase AOP2 |
| mi2118a | Solyc10g050115 | 3 | 5.699 | 891 | Transposon Ty3-I Gag-Pol polyprotein |
| mi2118a | Solyc11g011350 | 3 | 19.87 | 1381 | disease resistance protein (TIR-NBS-LRR class) |
| mi2118a | Solyc12g009450 | 3 | 19.542 | 727 | Disease resistance protein (CC-NBS-LRR class) family |
| mi2118a | Solyc12g056490 | 3 | 23.311 | 324 | WD40 repeat-containing protein |
| mi2118a | Solyc01g094520 | 3.5 | 20.797 | 978 | F-box/kelch-repeat protein |
| mi2118a | Solyc01g112260 | 3.5 | 30.089 | 316 | Phosphoenolpyruvate carboxylase |
| mi2118a | Solyc02g064680 | 3.5 | 19.849 | 1125 | Calcium-transporting ATPase |
| mi2118a | Solyc02g090860 | 3.5 | 19.613 | 1387 | Phenylalanyl-tRNA synthetase alpha chain |
| mi2118a | Solyc03g025190 | 3.5 | 21.744 | 803 | anthocyanin permease |
| mi2118a | Solyc03g112630 | 3.5 | 16.495 | 3497 | Sec14p-like phosphatidylinositol transfer family protein |
| mi2118a | Solyc04g011960 | 3.5 | 15.189 | 521 | Disease resistance protein (CC-NBS-LRR class) family |
| mi2118a | Solyc04g011980 | 3.5 | 16.151 | 521 | Disease resistance protein (CC-NBS-LRR class) family |
| mi2118a | Solyc04g011990 | 3.5 | 15.575 | 1077 | Disease resistance protein (NBS-LRR class) family |
| mi2118a | Solyc04g012000 | 3.5 | 17.228 | 221 | NBS-coding resistance gene analog |
| mi2118a | Solyc04g012010 | 3.5 | 18.466 | 541 | Disease resistance protein (NBS-LRR class) family |
| mi2118a | Solyc05g010240 | 3.5 | 11.485 | 3748 | Chaperonin-60 beta subunit |
| mi2118a | Solyc06g061215 | 3.5 | 20.712 | 170 | Proteinase inhibitor II |
| mi2118a | Solyc06g069390 | 3.5 | 24.527 | 1442 | D-aminoacyl-tRNA deacylase |
| mi2118a | Solyc07g063430 | 3.5 | 13.261 | 643 | Peroxisomal membrane (Mpv17/PMP22) family protein |
| mi2118a | Solyc08g066500 | 3.5 | 13.834 | 554 | Homeobox leucine-zipper protein |
| mi2118a | Solyc09g005290 | 3.5 | 20.812 | 668 | Nbs-lrr resistance protein |
| mi2118a | Solyc09g091990 | 3.5 | 18.213 | 665 | Kinase family protein |
| mi2118a | Solyc10g087013 | 3.5 | 18.136 | 838 | Cytochrome P450 |
| mi2118a | Solyc12g056960 | 3.5 | 13.791 | 484 | Glucan 1 |
| mi2118a | Solyc01g008800 | 4 | 14.87 | 1561 | disease resistance protein (TIR-NBS-LRR class) |
| mi2118a | Solyc01g111160 | 4 | 17.893 | 181 | far-red elongated hypocotyls 3 |
| mi2118a | Solyc02g081870 | 4 | 16.621 | 267 | Pleiotropic drug resistance ABC transporter |
| mi2118a | Solyc02g093340 | 4 | 15.426 | 1078 | RNA-binding (RRM/RBD/RNP motifs) family protein |
| mi2118a | Solyc03g111140 | 4 | 18.294 | 766 | Malate synthase |
| mi2118a | Solyc03g115740 | 4 | 10.662 | 2443 | Xyloglucan alpha-1 |
| mi2118a | Solyc05g008340 | 4 | 23.996 | 791 | Core-2/I-branching beta-1 |
| mi2118a | Solyc05g009470 | 4 | 14.478 | 2663 | Alpha-glucosidase |
| mi2118a | Solyc06g009533 | 4 | 20.01 | 1085 | Kinase family protein |
| mi2118a | Solyc06g065820 | 4 | 14.049 | 582 | Ethylene Response Factor H.1 |
| mi2118a | Solyc06g068700 | 4 | 21.249 | 2627 | Calreticulin/calnexin |
| mi2118a | Solyc07g008950 | 4 | 20.753 | 2523 | Methionyl-tRNA synthetase family protein |
| mi2118a | Solyc07g008955 | 4 | 20.753 | 2631 | Unknown protein |
| mi2118a | Solyc07g041030 | 4 | 22.468 | 277 | DNA topoisomerase |
| mi2118a | Solyc08g013900 | 4 | 19.494 | 4686 | Plant regulator RWP-RK family protein |
| mi2118a | Solyc08g082000 | 4 | 17.749 | 789 | Homeobox-leucine zipper HOX24 |
| mi2118a | Solyc09g007710 | 4 | 26.176 | 986 | Disease resistance protein (TIR-NBS-LRR class) family |
| mi2118a | Solyc09g075010 | 4 | 4.321 | 683 | HSP20-like chaperones superfamily protein |
| mi2118a | Solyc11g045350 | 4 | 19.839 | 3532 | Plant regulator RWP-RK family protein |
| mi2118a | Solyc11g062220 | 4 | 19.961 | 5265 | Zinc finger CCCH domain-containing protein 44 |
| mi2118b | Solyc06g005410 | 1 | 15.878 | 592 | TAS5 |
| mi2118b | Solyc06g005410 | 1 | 15.569 | 1639 | TAS5 |
| mi2118b | Solyc02g032650 | 2 | 15.284 | 796 | Disease resistance protein (TIR-NBS-LRR class) |
| mi2118b | Solyc01g105340 | 2.5 | 20.142 | 810 | Chaperone protein DnaJ |
| mi2118b | Solyc01g113620 | 2.5 | 16.895 | 806 | Disease resistance protein (TIR-NBS-LRR class) family |
| mi2118b | Solyc04g009110 | 2.5 | 18.119 | 572 | Nbs-*lrr* resistance protein |
| mi2118b | Solyc04g009130 | 2.5 | 20.332 | 584 | Nbs-*lrr* resistance protein |
| mi2118b | Solyc04g009290 | 2.5 | 18.898 | 572 | Disease resistance protein |
| mi2118b | Solyc04g026110 | 2.5 | 22.415 | 491 | Disease resistance family protein |
| mi2118b | Solyc05g009750 | 2.5 | 17.541 | 164 | NBS-LRR resistance protein-like protein |
| mi2118b | Solyc08g075630 | 2.5 | 26.52 | 743 | NBS-LRR resistance protein |
| mi2118b | Solyc08g076000 | 2.5 | 25.842 | 851 | NBS-LRR resistance protein |
| mi2118b | Solyc01g102920 | 3 | 14.713 | 635 | Disease resistance protein (TIR-NBS-LRR class) |
| mi2118b | Solyc02g030290 | 3 | 19.46 | 281 | Nbs-*lrr* resistance protein |
| mi2118b | Solyc02g037540 | 3 | 23.811 | 479 | Disease resistance protein |
| mi2118b | Solyc06g062440 | 3 | 22.729 | 592 | Disease resistance protein |
| mi2118b | Solyc08g075640 | 3 | 24.051 | 836 | NBS-LRR resistance protein |
| mi2118b | Solyc09g075010 | 3 | 16.92 | 611 | HSP20-like chaperones superfamily protein |
| mi2118b | Solyc09g098100 | 3 | 18.265 | 2059 | CC-NBS-LRR_Solyc09g098100 |
| mi2118b | Solyc01g110000 | 3.5 | 16.414 | 2336 | Beta-galactosidase |
| mi2118b | Solyc02g036270 | 3.5 | 16.852 | 679 | Disease resistance protein (NBS-LRR class) family |
| mi2118b | Solyc03g123630 | 3.5 | 23.333 | 1670 | pectin methylesterase |
| mi2118b | Solyc04g009120 | 3.5 | 22.379 | 599 | Nbs-*lrr* resistance protein |
| mi2118b | Solyc04g025820 | 3.5 | 20.87 | 312 | Disease resistance protein |
| mi2118b | Solyc04g025840 | 3.5 | 20.843 | 491 | Disease resistance family protein |
| mi2118b | Solyc05g008070 | 3.5 | 21.429 | 530 | Disease resistance protein |
| mi2118b | Solyc05g014030 | 3.5 | 21.401 | 738 | Regulator of chromosome condensation (RCC1) family protein |
| mi2118b | Solyc06g007780 | 3.5 | 21.021 | 2776 | Nuclear transport factor 2 (NTF2) |
| mi2118b | Solyc07g039400 | 3.5 | 22.502 | 491 | Disease resistance protein |
| mi2118b | Solyc07g039420 | 3.5 | 22.458 | 566 | Disease resistance protein (NBS-LRR class) family |
| mi2118b | Solyc08g067060 | 3.5 | 24.14 | 674 | Pentatricopeptide repeat superfamily protein |
| mi2118b | Solyc08g074250 | 3.5 | 15.378 | 545 | Disease resistance protein (CC-NBS-LRR class) family |
| mi2118b | Solyc11g008140 | 3.5 | 11.101 | 1224 | Pectate lyase |
| mi2118b | Solyc12g006040 | 3.5 | 21.275 | 662 | NBS-LRR protein |
| mi2118b | Solyc01g105775 | 4 | 23.141 | 221 | Carbonic anhydrase |
| mi2118b | Solyc01g111100 | 4 | 18.433 | 635 | Neutral invertase |
| mi2118b | Solyc02g079310 | 4 | 16.167 | 387 | Equilibrative nucleoside transporter family protein |
| mi2118b | Solyc02g079350 | 4 | 14.312 | 2490 | Equilibrative nucleoside transporter family protein |
| mi2118b | Solyc02g083960 | 4 | 19.892 | 3334 | 2-oxoglutarate and Fe-dependent oxygenase-like protein |
| mi2118b | Solyc03g007330 | 4 | 22.962 | 983 | ATP-dependent zinc metalloprotease FTSH protein |
| mi2118b | Solyc03g083430 | 4 | 21.732 | 1551 | Splicing factor 3A subunit 3 |
| mi2118b | Solyc04g005540 | 4 | 17.383 | 850 | Disease resistance protein (NBS-LRR class) family |
| mi2118b | Solyc04g005550 | 4 | 21.364 | 868 | Disease resistance protein (NBS-LRR class) family |
| mi2118b | Solyc04g071260 | 4 | 21.507 | 2316 | Actin |
| mi2118b | Solyc05g014760 | 4 | 21.583 | 3126 | Kinase family protein |
| mi2118b | Solyc05g018720 | 4 | 17.342 | 62 | NBS-coding resistance protein |
| mi2118b | Solyc07g005770 | 4 | 24.094 | 539 | Disease resistance protein |
| mi2118b | Solyc07g064700 | 4 | 25.463 | 1362 | Bromodomain-containing protein |
| mi2118b | Solyc09g005120 | 4 | 18.535 | 483 | DnaJ domain-containing protein |
| mi2118b | Solyc09g076010 | 4 | 8.678 | 370 | Acyl-CoA N-acyltransferase |
| mi2118b | Solyc10g008230 | 4 | 19.11 | 548 | Disease resistance protein |
| mi2118b | Solyc11g069020 | 4 | 20.947 | 497 | Disease resistance protein |
| mi2118b | Solyc12g017800 | 4 | 23.624 | 1102 | NBS-LRR class disease resistance protein |
| mi2118b | Solyc12g099060 | 4 | 23.521 | 644 | Disease resistance protein |
| mi2118b | Solyc12g099940 | 4 | 15.941 | 2435 | Acyl-CoA N-acyltransferase |
| miR482b | Solyc04g009070 | 0 | 25.758 | 211 | Disease resistance family protein |
| miR482b | Solyc02g036270 | 1.5 | 12.267 | 681 | Disease resistance protein (NBS-LRR class) family |
| miR482b | Solyc04g009120 | 1.5 | 21.183 | 601 | Nbs-lrr resistance protein |
| miR482b | Solyc05g008070 | 1.5 | 21.303 | 532 | Disease resistance protein |
| miR482b | Solyc04g025820 | 2 | 19.14 | 314 | Disease resistance protein |
| miR482b | Solyc04g025840 | 2 | 19.226 | 493 | Disease resistance family protein |
| miR482b | Solyc07g039420 | 2 | 23.494 | 568 | Disease resistance protein (NBS-LRR class) family |
| miR482b | Solyc11g065780 | 2 | 21.191 | 786 | Disease resistance protein |
| miR482b | Solyc12g017800 | 2 | 24.453 | 1104 | NBS-LRR class disease resistance protein |
| miR482b | Solyc01g067165 | 2.5 | 20.018 | 529 | Disease resistance protein (CC-NBS-LRR class) family protein |
| miR482b | Solyc04g009130 | 2.5 | 20.794 | 586 | Nbs-lrr resistance protein |
| miR482b | Solyc04g009290 | 2.5 | 23.41 | 574 | Disease resistance protein |
| miR482b | Solyc07g039400 | 2.5 | 22.497 | 493 | Disease resistance protein |
| miR482b | Solyc10g054970 | 2.5 | 19.563 | 538 | CCNBS gene |
| miR482b | Solyc10g054990 | 2.5 | 16.38 | 520 | Disease resistance protein (NBS-LRR class) family |
| miR482b | Solyc10g055170 | 2.5 | 21.742 | 46 | Disease resistance protein (CC-NBS-LRR class) family protein |
| miR482b | Solyc11g006530 | 2.5 | 18.804 | 526 | Disease resistance protein |
| miR482b | Solyc11g006630 | 2.5 | 21.57 | 532 | Disease resistance protein |
| miR482b | Solyc04g009240 | 3 | 27.611 | 565 | Nbs-lrr resistance protein |
| miR482b | Solyc04g009250 | 3 | 28.375 | 577 | Nbs-lrr resistance protein |
| miR482b | Solyc04g009660 | 3 | 28.833 | 550 | Nbs-lrr resistance protein |
| miR482b | Solyc04g009690 | 3 | 27.03 | 562 | Nbs-lrr resistance protein |
| miR482b | Solyc07g005770 | 3 | 23.779 | 541 | Disease resistance protein |
| miR482b | Solyc08g005440 | 3 | 21.814 | 627 | NBS-LRR disease resistance protein |
| miR482b | Solyc10g051050 | 3 | 21.243 | 929 | Disease resistance protein |
| miR482b | Solyc11g020090 | 3 | 20.197 | 103 | Disease resistance protein |
| miR482b | Solyc11g020100 | 3 | 23.736 | 1110 | Disease resistance protein |
| miR482b | Solyc11g069925 | 3 | 24.835 | 622 | Disease resistance protein |
| miR482b | Solyc01g113620 | 3.5 | 16.593 | 808 | Disease resistance protein (TIR-NBS-LRR class) family |
| miR482b | Solyc02g032650 | 3.5 | 15.288 | 798 | Disease resistance protein (TIR-NBS-LRR class) |
| miR482b | Solyc02g070730 | 3.5 | 25.498 | 412 | NBS-LRR resistance protein |
| miR482b | Solyc04g005540 | 3.5 | 15.547 | 852 | Disease resistance protein (NBS-LRR class) family |
| miR482b | Solyc04g005550 | 3.5 | 18.848 | 870 | Disease resistance protein (NBS-LRR class) family |
| miR482b | Solyc04g009110 | 3.5 | 17.68 | 574 | Nbs-lrr resistance protein |
| miR482b | Solyc04g056746 | 3.5 | 16.29 | 687 | Pentatricopeptide repeat-containing protein |
| miR482b | Solyc05g006630 | 3.5 | 29.979 | 1034 | disease resistance protein (TIR-NBS-LRR class) |
| miR482b | Solyc05g007170 | 3.5 | 22.989 | 5882 | Disease resistance protein |
| miR482b | Solyc06g072000 | 3.5 | 20.074 | 307 | P-loop containing nucleoside triphosphate hydrolases superfamily protein |
| miR482b | Solyc07g044790 | 3.5 | 21.329 | 1223 | Pvr4 |
| miR482b | Solyc07g044797 | 3.5 | 21.329 | 286 | CC-NBS-LRR disease resistance protein |
| miR482b | Solyc07g049700 | 3.5 | 24.47 | 544 | Disease resistance protein |
| miR482b | Solyc09g065560 | 3.5 | 14.452 | 1794 | Sulfate transporter |
| miR482b | Solyc11g006520 | 3.5 | 23.607 | 819 | Disease resistance protein |
| miR482b | Solyc11g006640 | 3.5 | 26.45 | 532 | Disease resistance protein |
| miR482b | Solyc11g068360 | 3.5 | 22.83 | 622 | Disease resistance protein |
| miR482b | Solyc11g069620 | 3.5 | 21.64 | 706 | Disease resistance protein |
| miR482b | Solyc12g044180 | 3.5 | 19.352 | 526 | CC-NBS-LRR disease resistance protein |
| miR482b | Solyc12g044190 | 3.5 | 18.858 | 526 | Disease resistance protein (CC-NBS-LRR class) family protein |
| miR482b | Solyc12g044200 | 3.5 | 18.45 | 1186 | Disease resistance protein (CC-NBS-LRR class) family protein |
| miR482b | Solyc02g037540 | 4 | 23.271 | 481 | Disease resistance protein |
| miR482b | Solyc03g078300 | 4 | 20.201 | 726 | Disease resistance protein |
| miR482b | Solyc04g009150 | 4 | 25.145 | 565 | Nbs-lrr resistance protein |
| miR482b | Solyc04g026110 | 4 | 21.107 | 493 | Disease resistance family protein |
| miR482b | Solyc04g048920 | 4 | 15.927 | 46 | CC-NBS-LRR disease resistance protein |
| miR482b | Solyc05g012740 | 4 | 20.929 | 1585 | Disease resistance protein |
| miR482b | Solyc07g053010 | 4 | 28.026 | 1171 | NBS-LRR type disease resistance protein |
| miR482b | Solyc10g045050 | 4 | 14.193 | 268 | Enolase |
| miR482b | Solyc10g051170 | 4 | 20.714 | 673 | Disease resistance protein |
| miR482b | Solyc10g055050 | 4 | 17.252 | 379 | CC-NBS-LRR disease resistance protein |
| miR482b | Solyc11g069990 | 4 | 23.911 | 1218 | I2C5 |
| miR482b | Solyc11g070020 | 4 | 23.911 | 1990 | Disease resistance protein |
| miR482b | Solyc11g071423 | 4 | 25.798 | 229 | Disease resistance protein |
| miR482b | Solyc11g071995 | 4 | 26.508 | 622 | Disease resistance protein |
| miR482b | Solyc12g006040 | 4 | 20.75 | 664 | NBS-LRR protein |
| miR482c | Solyc08g075630 | 1.5 | 24.851 | 745 | NBS-LRR resistance protein |
| miR482c | Solyc08g076000 | 1.5 | 24.722 | 853 | NBS-LRR resistance protein |
| miR482c | Solyc02g021140 | 2.5 | 7.72 | 4 | Superoxide dismutase |
| miR482c | Solyc02g078280 | 3 | 11.484 | 1597 | DNA ligase-like protein |
| miR482c | Solyc05g006630 | 3 | 29.979 | 1034 | disease resistance protein (TIR-NBS-LRR class) |
| miR482c | Solyc06g076350 | 3 | 15.598 | 392 | LePCL1 |
| miR482c | Solyc11g011560 | 3 | 19.304 | 1085 | PHD finger protein family |
| miR482c | Solyc11g065780 | 3 | 21.191 | 786 | Disease resistance protein |
| miR482c | Solyc00g006530 | 3.5 | 25.591 | 771 | Calmodulin-binding protein |
| miR482c | Solyc01g102660 | 3.5 | 18.543 | 218 | Glutathione S-transferase |
| miR482c | Solyc02g032650 | 3.5 | 15.288 | 798 | Disease resistance protein (TIR-NBS-LRR class) |
| miR482c | Solyc02g036270 | 3.5 | 12.267 | 681 | Disease resistance protein (NBS-LRR class) family |
| miR482c | Solyc04g005540 | 3.5 | 15.547 | 852 | Disease resistance protein (NBS-LRR class) family |
| miR482c | Solyc04g005550 | 3.5 | 18.848 | 870 | Disease resistance protein (NBS-LRR class) family |
| miR482c | Solyc04g009110 | 3.5 | 17.68 | 574 | Nbs-lrr resistance protein |
| miR482c | Solyc04g009150 | 3.5 | 25.145 | 565 | Nbs-lrr resistance protein |
| miR482c | Solyc04g025160 | 3.5 | 19.754 | 340 | ATPase |
| miR482c | Solyc04g026110 | 3.5 | 21.107 | 493 | Disease resistance family protein |
| miR482c | Solyc04g080590 | 3.5 | 25.393 | 748 | Proteasome subunit alpha type |
| miR482c | Solyc06g060360 | 3.5 | 21.646 | 1010 | Adenine nucleotide alpha hydrolases-like superfamily protein |
| miR482c | Solyc06g083875 | 3.5 | 17.908 | 247 | pollen Ole e I family allergen protein |
| miR482c | Solyc07g053200 | 3.5 | 12.314 | 923 | Adenine nucleotide alpha hydrolases-like superfamily protein |
| miR482c | Solyc08g065740 | 3.5 | 23.005 | 475 | Vacuolar processing enzyme |
| miR482c | Solyc08g068040 | 3.5 | 19.304 | 2071 | zinc finger FYVE domain protein |
| miR482c | Solyc10g007200 | 3.5 | 11.289 | 503 | Hexosyltransferase |
| miR482c | Solyc10g076440 | 3.5 | 18.053 | 475 | Disease resistance protein |
| miR482c | Solyc11g017370 | 3.5 | 19.701 | 1910 | Pentatricopeptide repeat-containing protein |
| miR482c | Solyc11g062010 | 3.5 | 20.533 | 9655 | Chromatin structure-remodeling complex subunit snf21 |
| miR482c | Solyc12g005230 | 3.5 | 13.933 | 4336 | Breast carcinoma-amplified sequence 3 |
| miR482c | Solyc12g019144 | 3.5 | 23.587 | 1538 | RING/U-box superfamily protein |
| miR482c | Solyc01g097390 | 4 | 20.058 | 1141 | NAD(P)-linked oxidoreductase superfamily protein |
| miR482c | Solyc01g103450 | 4 | 30.773 | 415 | Heat shock protein 70 |
| miR482c | Solyc02g005180 | 4 | 25.396 | 273 | Sugar facilitator protein 2 |
| miR482c | Solyc02g070730 | 4 | 25.498 | 412 | NBS-LRR resistance protein |
| miR482c | Solyc02g078790 | 4 | 14.982 | 1601 | Transcription factor jumonji domain protein |
| miR482c | Solyc02g080960 | 4 | 18.235 | 39 | transmembrane protein |
| miR482c | Solyc03g097980 | 4 | 17.739 | 550 | Guanine nucleotide-binding alpha-2 subunit |
| miR482c | Solyc03g113620 | 4 | 28.565 | 230 | MYB transcription factor |
| miR482c | Solyc05g005460 | 4 | 23.644 | 2043 | Quinone oxidoreductase-like protein |
| miR482c | Solyc05g018370 | 4 | 5.408 | 808 | Leguminosin group485 secreted peptide |
| miR482c | Solyc06g062440 | 4 | 22.498 | 595 | Disease resistance protein |
| miR482c | Solyc06g068210 | 4 | 6.629 | 2252 | Protein FAR1-RELATED SEQUENCE 8 |
| miR482c | Solyc07g052760 | 4 | 16.463 | 540 | DNA-binding storekeeper protein-related transcriptional regulator |
| miR482c | Solyc07g055380 | 4 | 19.526 | 1969 | Disease resistance protein (TIR-NBS-LRR class) |
| miR482c | Solyc07g055610 | 4 | 20.758 | 553 | Disease resistance protein (TIR-NBS-LRR class) |
| miR482c | Solyc09g007830 | 4 | 14.32 | 265 | Cytokinin riboside 5'-monophosphate phosphoribohydrolase |
| miR482c | Solyc11g010660 | 4 | 12.438 | 988 | protein SGT1 |
| miR482c | Solyc11g062010 | 4 | 21.236 | 9856 | Chromatin structure-remodeling complex subunit snf21 |
| miR482c | Solyc11g069830 | 4 | 21.579 | 309 | ATPase ASNA1 |
| miR482e | Solyc11g006530 | 1.5 | 18.804 | 526 | Disease resistance protein |
| miR482e | Solyc11g006630 | 1.5 | 21.57 | 532 | Disease resistance protein |
| miR482e | Solyc07g049700 | 2 | 24.47 | 544 | Disease resistance protein |
| miR482e | Solyc01g067165 | 2.5 | 20.018 | 529 | Disease resistance protein (CC-NBS-LRR class) family protein |
| miR482e | Solyc04g009070 | 2.5 | 25.758 | 211 | Disease resistance family protein |
| miR482e | Solyc05g008070 | 2.5 | 21.303 | 532 | Disease resistance protein |
| miR482e | Solyc06g074760 | 2.5 | 15.026 | 1529 | Ring/U-Box superfamily protein |
| miR482e | Solyc10g054970 | 2.5 | 19.563 | 538 | CCNBS gene |
| miR482e | Solyc10g054990 | 2.5 | 16.38 | 520 | Disease resistance protein (NBS-LRR class) family |
| miR482e | Solyc10g055170 | 2.5 | 21.742 | 46 | Disease resistance protein (CC-NBS-LRR class) family protein |
| miR482e | Solyc11g020100 | 2.5 | 23.736 | 1110 | Disease resistance protein |
| miR482e | Solyc12g009450 | 2.5 | 19.509 | 729 | Disease resistance protein (CC-NBS-LRR class) family |
| miR482e | Solyc12g017800 | 2.5 | 24.453 | 1104 | NBS-LRR class disease resistance protein |
| miR482e | Solyc01g014840 | 3 | 16.059 | 649 | disease resistance protein (TIR-NBS-LRR class) |
| miR482e | Solyc01g108460 | 3 | 19.975 | 5259 | Carboxypeptidase |
| miR482e | Solyc04g009250 | 3 | 28.375 | 577 | Nbs-lrr resistance protein |
| miR482e | Solyc04g009660 | 3 | 28.833 | 550 | Nbs-lrr resistance protein |
| miR482e | Solyc04g009690 | 3 | 27.03 | 562 | Nbs-lrr resistance protein |
| miR482e | Solyc05g032850 | 3 | 17.25 | 2848 | evolutionarily conserved C-terminal region 2 |
| miR482e | Solyc07g027020 | 3 | 15.263 | 914 | Protein kinase family protein |
| miR482e | Solyc09g091050 | 3 | 15.779 | 833 | Calcium-dependent lipid-binding (CaLB domain) family protein |
| miR482e | Solyc11g006520 | 3 | 23.607 | 819 | Disease resistance protein |
| miR482e | Solyc11g006640 | 3 | 26.45 | 532 | Disease resistance protein |
| miR482e | Solyc11g069925 | 3 | 24.835 | 622 | Disease resistance protein |
| miR482e | Solyc12g006040 | 3 | 20.75 | 664 | NBS-LRR protein |
| miR482e | Solyc01g008800 | 3.5 | 16.457 | 1564 | disease resistance protein (TIR-NBS-LRR class) |
| miR482e | Solyc01g066020 | 3.5 | 26.023 | 658 | disease resistance protein (TIR-NBS-LRR class) |
| miR482e | Solyc02g032200 | 3.5 | 18.488 | 769 | Disease resistance protein (TIR-NBS-LRR class) family |
| miR482e | Solyc02g036270 | 3.5 | 12.267 | 681 | Disease resistance protein (NBS-LRR class) family |
| miR482e | Solyc03g046207 | 3.5 | 24.478 | 703 | Disease resistance protein (CC-NBS-LRR class) family protein |
| miR482e | Solyc04g005540 | 3.5 | 15.547 | 852 | Disease resistance protein (NBS-LRR class) family |
| miR482e | Solyc04g005550 | 3.5 | 18.848 | 870 | Disease resistance protein (NBS-LRR class) family |
| miR482e | Solyc04g009120 | 3.5 | 21.183 | 601 | Nbs-lrr resistance protein |
| miR482e | Solyc04g074865 | 3.5 | 16.828 | 1400 | Retrovirus-related Pol polyprotein from transposon TNT 1-94 |
| miR482e | Solyc05g006630 | 3.5 | 29.979 | 1034 | disease resistance protein (TIR-NBS-LRR class) |
| miR482e | Solyc07g005770 | 3.5 | 23.779 | 541 | Disease resistance protein |
| miR482e | Solyc07g044790 | 3.5 | 21.329 | 1223 | Pvr4 |
| miR482e | Solyc07g044797 | 3.5 | 21.329 | 286 | CC-NBS-LRR disease resistance protein |
| miR482e | Solyc10g051050 | 3.5 | 21.243 | 928 | Disease resistance protein |
| miR482e | Solyc11g011350 | 3.5 | 20.579 | 1383 | disease resistance protein (TIR-NBS-LRR class) |
| miR482e | Solyc11g020090 | 3.5 | 20.197 | 103 | Disease resistance protein |
| miR482e | Solyc11g068360 | 3.5 | 22.83 | 622 | Disease resistance protein |
| miR482e | Solyc11g069300 | 3.5 | 25.111 | 2436 | Kinase family protein |
| miR482e | Solyc11g069620 | 3.5 | 21.64 | 706 | Disease resistance protein |
| miR482e | Solyc11g069990 | 3.5 | 23.911 | 1218 | I2C5 |
| miR482e | Solyc11g070020 | 3.5 | 23.911 | 1990 | Disease resistance protein |
| miR482e | Solyc11g071423 | 3.5 | 25.798 | 229 | Disease resistance protein |
| miR482e | Solyc11g071995 | 3.5 | 26.508 | 622 | Disease resistance protein |
| miR482e | Solyc12g005970 | 3.5 | 15.572 | 493 | Disease resistance protein (CC-NBS-LRR class) family |
| miR482e | Solyc12g044180 | 3.5 | 19.352 | 526 | CC-NBS-LRR disease resistance protein |
| miR482e | Solyc12g044190 | 3.5 | 18.858 | 526 | Disease resistance protein (CC-NBS-LRR class) family protein |
| miR482e | Solyc12g044200 | 3.5 | 18.45 | 1186 | Disease resistance protein (CC-NBS-LRR class) family protein |
| miR482e | Solyc01g087200 | 4 | 19.546 | 517 | Disease resistance protein |
| miR482e | Solyc01g100310 | 4 | 14.841 | 947 | Calmodulin-binding protein |
| miR482e | Solyc02g032650 | 4 | 15.288 | 798 | Disease resistance protein (TIR-NBS-LRR class) |
| miR482e | Solyc02g073574 | 4 | 17.186 | 583 | Disease resistance protein |
| miR482e | Solyc02g084450 | 4 | 17.186 | 2110 | Disease resistance protein |
| miR482e | Solyc03g078300 | 4 | 20.201 | 727 | Disease resistance protein |
| miR482e | Solyc04g011590 | 4 | 30.314 | 622 | Amino acid transporter |
| miR482e | Solyc04g011960 | 4 | 15.415 | 523 | Disease resistance protein (CC-NBS-LRR class) family |
| miR482e | Solyc04g011980 | 4 | 16.137 | 523 | Disease resistance protein (CC-NBS-LRR class) family |
| miR482e | Solyc04g011990 | 4 | 15.741 | 1079 | Disease resistance protein (NBS-LRR class) family |
| miR482e | Solyc04g012000 | 4 | 17.872 | 223 | NBS-coding resistance gene analog |
| miR482e | Solyc04g012010 | 4 | 18.376 | 543 | Disease resistance protein (NBS-LRR class) family |
| miR482e | Solyc04g048920 | 4 | 15.927 | 46 | CC-NBS-LRR disease resistance protein |
| miR482e | Solyc08g079150 | 4 | 16.89 | 22 | SAUR-like auxin-responsive protein family |
| miR482e | Solyc09g007710 | 4 | 25.33 | 989 | Disease resistance protein (TIR-NBS-LRR class) family |
| miR482e | Solyc10g055050 | 4 | 17.252 | 379 | CC-NBS-LRR disease resistance protein |
| miR482e | Solyc11g065780 | 4 | 21.191 | 786 | Disease resistance protein |
| miR482e | Solyc12g005520 | 4 | 23.107 | 121 | Disease resistance protein (CC-NBS-LRR class) family |
| miR482e | Solyc12g096920 | 4 | 17.72 | 544 | Disease resistance protein (CC-NBS-LRR class) family protein |

**Table S3.** Target prediction for all siRNA-producing NLRs. Summary of sNLs, with their gene id, class of NLR protein based on the phylogen genetic analysis of a previous study (38) and not in the presence of representative domains, total counts for 21-nt sRNAs (nRPM), and targeting scores for each individual microRNA (red indicates stronger targeting prediction). *TAS5* (bottom) is added for reference.

| General information |  |  | Target Prediction |  |  |  |  |
| --- | --- | --- | --- | --- | --- | --- | --- |
| Gene_ID | NLR class | sRNA prod. | miR482b | miR482e | miR482c | miR2118a | miR2118b |
| Solyc11g065780 | CNL | 1159.4 | 2 | 3 | 4 | - | - |
| Solyc05g008070 | CNL | 569.4 | 1.5 | 2.5 | - | - | 3.5 |
| Solyc02g036270 | CNL - LRR1 | 514.6 | 1.5 | 3.5 | 3.5 | - | 3.5 |
| Solyc04g005540 | CNL - LRR2 | 303.2 | 3.5 | 3.5 | 3.5 | - | 4 |
| Solyc11g071995 | CNL | 259.4 | 4 | 3.5 | - | - | - |
| Solyc04g005550 | CNL | 129.6 | 3.5 | 3.5 | 3.5 | - | 4 |
| Solyc10g051050 | CNL | 128.0 | 3 | 3.5 | - | - | - |
| Solyc09g064610 | CNL | 77.7 | - | - | - | - | - |
| Solyc01g008800 | TNL | 74.7 | - | 3.5 | - | 4 | - |
| Solyc08g007630 | CNL | 70.0 | - | - | - | - | - |
| Solyc05g009630 | CNL | 66.9 | - | - | - | - | - |
| Solyc11g069990 | CNL | 65.3 | 4 | 3.5 | - | - | - |
| Solyc11g069620 | CNL | 64.9 | 3.5 | 3.5 | - | - | - |
| Solyc11g069925 | CNL | 58.4 | 3 | 3 | - | - | - |
| Solyc05g005330 | CNL | 48.2 | - | - | - | - | - |
| Solyc11g068360 | CNL | 34.4 | 3.5 | 3.5 | - | - | - |
| Solyc07g049700 | CNL | 25.0 | 3.5 | 2 | - | - | - |
| Solyc11g071410 | CNL | 23.3 | - | - | - | - | - |
| Solyc10g085460 | CNL | 19.9 | - | - | - | - | - |
| Solyc11g011350 | TNL | 18.2 | - | 3.5 | - | 3 | - |
| Solyc11g006640 | CNL | 17.8 | 3.5 | 3 | - | - | - |
| Solyc12g044190 | CNL | 17.3 | 3.5 | 3.5 | - | - | - |
| Solyc11g020100 | CNL | 17.3 | 3 | 2.5 | - | - | - |
| Solyc12g044200 | CNL | 17.2 | 3.5 | 3.5 | - | - | - |
| Solyc07g005770 | CNL | 15.5 | 3 | 3.5 | - | - | 4 |
| Solyc09g018220 | CNL | 14.7 | - | - | - | - | - |
| Solyc11g064770 | CNL | 14.3 | - | - | - | - | - |
| Solyc02g032650 | TNL | 14.3 | 3.5 | 4 | 3.5 | - | 2 |
| Solyc12g006040 | CNL | 13.7 | 4 | 3 | - | - | 3.5 |
| Solyc08g076000 | CNL | 12.5 | - | - | 1.5 | - | 2.5 |
| Solyc11g069660 | CNL | 12.0 | - | - | - | - | - |
| Solyc09g098130 | CNL | 11.1 | - | - | - | - | - |
| Solyc06g005410 | TAS5 | 431.2 | - | - | - | - | 1 / 2 |

**Table S4.** Summary of degradome (PARE) signatures. Summary of validated degradome products for miR482/2118 members on the whole tomato transcriptome (p-value < 0.05). Peak category refers to PARE read abundances of that position correspond to (0) > 90th percentile, (1) > 75th, (2) > 50th percentile, (3) < 50th percentile of total PARE read signatures in the genome.

| miRname | Target | Score | Cleavage Position | Reads | Proportion of reads in 10nt window | Peak category | Corrected p-value | Annotation |
| --- | --- | --- | --- | --- | --- | --- | --- | --- |
| miR2118b/a | Solyc06g005410 | 1 / 4 | 513 | 62 | 1 | 0 | 0.0001/0.0058 | TAS5 |
| miR482b/c | Solyc02g036270 | 1.5 / 3 | 1017 | 12 | 1 | 2 | 0.0058/ 0.0155 | LRR1 |
| miR2118b | Solyc06g005410 | 2 | 2536 | 18 | 0.3 | 2 | 0.0061 | TAS5 |
| miR482c | Solyc04g005540 | 3 | 864 | 7 | 1 | 2 | 0.0155 | LRR2 |
| miR482e | Solyc11g013750 | 4 | 1778 | 9 | 1 | 2 | 0.0401 | Disease resistance protein (NLR class) family |
| miR2118a | Solyc06g064550 | 7 | 6499 | 8 | 1 | 2 | 0.042 | Aspartokinase-homoserine dehydrogenase |
| miR482c | Solyc06g062440 | 3.5 | 607 | 7 | 1 | 2 | 0.0475 | Disease resistance protein |
| miR2118a | Solyc08g065220 | 7 | 4795 | 4 | 1 | 3 | 0.0487 | Glycine decarboxylase p-protein |

**Table S5.** Summary of phasing signatures. Top 20 phasing signatures in our sRNA dataset mapping to tomato genes. Colour code indicates when the gene is a predicted to be a preferential target of (blue) miR482 or (red) miR2118.

| Gene ID | Start | End | phaseR score | Annotation |
| --- | --- | --- | --- | --- |
| Solyc02g036270 | 278 | 3514 | -26.9 | LRR1 |
| Solyc06g005410 | 361 | 1513 | -24.6 | TAS5 |
| Solyc10g051050 | 938 | 2624 | -22.9 | Disease resistance protein (NLR class) family |
| Solyc09g074520 | 409 | 2264 | -21.0 | Transport inhibitor response 1 (TIR1) |
| Solyc05g008070 | 287 | 2479 | -18.8 | Disease resistance protein (NLR class) family |
| Solyc04g005540 | 431 | 3965 | -18.3 | LRR2 |
| Solyc05g009630 | 701 | 2948 | -17.4 | Disease resistance protein (NLR class) family |
| Solyc11g069990 | 763 | 3397 | -16.5 | Disease resistance protein (NLR class) family |
| Solyc11g065820 | 2293 | 6253 | -16.0 | Disease resistance protein (NLR class) family |
| Solyc12g099870 | 1691 | 2072 | -15.5 | LRR RLK |
| Solyc11g011350 | 780 | 3629 | -15.5 | Disease resistance protein (NLR class) family |
| Solyc06g048960 | 518 | 3675 | -15.3 | Dicer-like 2a (DCL2a) |
| Solyc05g051230 | 2225 | 2876 | -14.2 | MOCS3-like |
| Solyc04g051190 | 154 | 1809 | -14.2 | P450 carotenoid β-hydrolase (CYP97A29) |
| Solyc11g071995 | 506 | 3586 | -14.1 | Disease resistance protein (NLR class) family |
| Solyc11g020100 | 957 | 2602 | -14.0 | Disease resistance protein (NLR class) family |
| Solyc01g058100 | 0 | 126 | -13.9 | NAD(P)H-quinone oxidoreductase subunit K |
| Solyc09g018220 | 562 | 2549 | -13.5 | Disease resistance protein (NLR class) family |
| Solyc02g032650 | 376 | 3091 | -13.3 | Disease resistance protein (NLR class) family |
| Solyc11g065790 | 12 | 409 | -13.2 | Disease resistance protein (NLR class) family |

**Table S6.** Most significant differential sRNA loci in STTM lines. Genetic loci with differential accumulation of sRNAs (any size-class), with their gene id, annotation, log2 fold changes, direction of the change, and adjusted p-value (cut-off of 0.05). Colour code indicates when the gene is a predicted to be a preferential target of (blue) miR482 or (red) miR2118. **Differential sRNA loci between STTM482.1 and wild type** Differential sRNA loci between MIM2118b and wild type

**Differential sRNA loci between STTM482.1 and wild type**
| Gene ID | Annotation | Log2(FC) | Condition | adj. p-value |
| --- | --- | --- | --- | --- |
| Solyc11g065780 | Disease resistance protein (NLR class) family | 1.596 | WT > STTM482.1 | 2.3E-06 |
| Solyc09g064610 | Disease resistance protein (NLR class) family | 3.255 | WT > STTM482.1 | 3.2E-05 |
| Solyc01g008790 | Non specific phospholipase C | 2.730 | WT > STTM482.1 | 2.3E-04 |
| Solyc01g067165 | Disease resistance protein (NLR class) family | 1.971 | WT > STTM482.1 | 3.4E-04 |
| Solyc01g008800 | Disease resistance protein (NLR class) family | 2.508 | WT > STTM482.1 | 4.5E-04 |
| Solyc04g017620 | F-box family protein | 2.065 | WT > STTM482.1 | 5.4E-04 |
| Solyc12g044190 | Disease resistance protein (NLR class) family | 1.924 | WT > STTM482.1 | 9.4E-04 |
| Solyc07g005770 | Disease resistance protein (NLR class) family | 2.787 | WT > STTM482.1 | 1.4E-03 |
| Solyc08g076000 | Disease resistance protein (NLR class) family | 2.336 | WT > STTM482.1 | 1.8E-03 |
| Solyc02g036270 | Disease resistance protein (NLR class) family | 1.921 | WT > STTM482.1 | 2.3E-03 |
| Solyc02g032650 | Disease resistance protein (NLR class) family | 2.140 | WT > STTM482.1 | 3.6E-03 |
| Solyc04g005550 | Disease resistance protein (NLR class) family | 2.306 | WT > STTM482.1 | 7.1E-03 |
| Solyc04g005540 | Disease resistance protein (NLR class) family | 1.751 | WT > STTM482.1 | 1.7E-02 |
| Solyc01g067147 | Asterix-like protein | 2.230 | WT > STTM482.1 | 3.2E-02 |
| Solyc01g100380 | Calreticulin | 1.322 | WT < STTM482.1 | 1.2E-02 |
| Solyc03g112330 | U-box domain-containing kinase family protein | 1.684 | WT < STTM482.1 | 2.5E-02 |
| Solyc03g112335 | O-acyltransferase (WSD1-like) family protein | 1.637 | WT < STTM482.1 | 4.0E-02 |
| Solyc09g097780 | Glycine-rich protein | 1.499 | WT < STTM482.1 | 4.8E-02 |

**Differential sRNA loci between MIM2118b and wild type**
| Gene ID | Annotation | Log2(FC) | Condition | adj. p-value |
| --- | --- | --- | --- | --- |
| Solyc06g005410 | TAS5 | 2.168 | WT > STTM2118b.5 | 8.09E-05 |

**Table S7.**
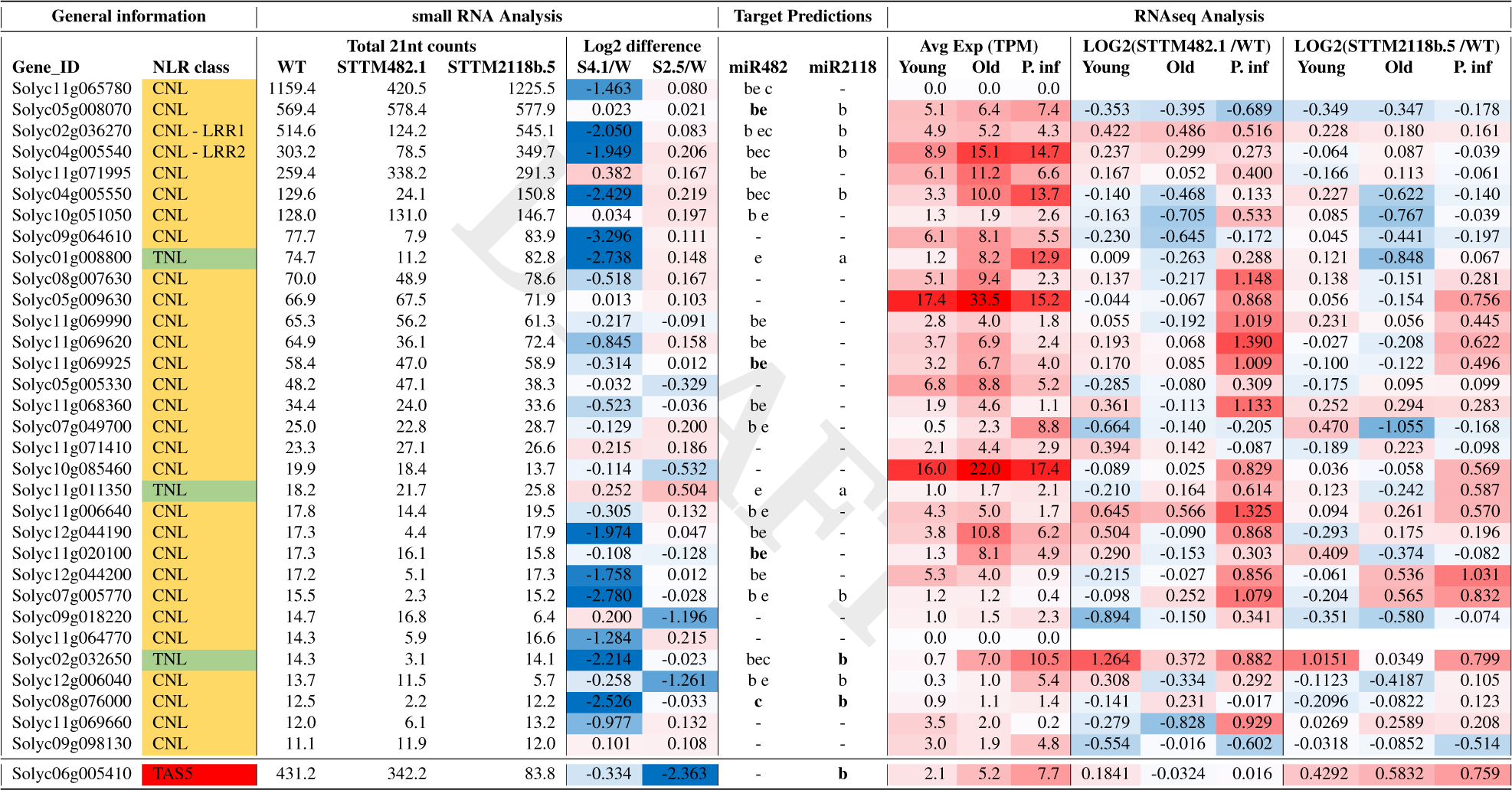
Summary of sNLs. Gene id, class of NLR protein based on the phylogenetic analysis of Andolfo et al. [2014]. Total counts for 21-nt sRNAs (nRPM) in wild type (WT), STTM482.1 and STTM2118b.5 lines. Log2 fold changes between WT and STTM lines (intensity of colour indicates stronger reduction). Summary of target prediction, with letters indicating the predicted targeting miRNA. Summary of RNAseq abundances in transcript per million (TPM) and Log2 fold changes between STTM lines and WT across conditions. SlyTAS5 (bottom) is added for additional reference.

**Table S8.**
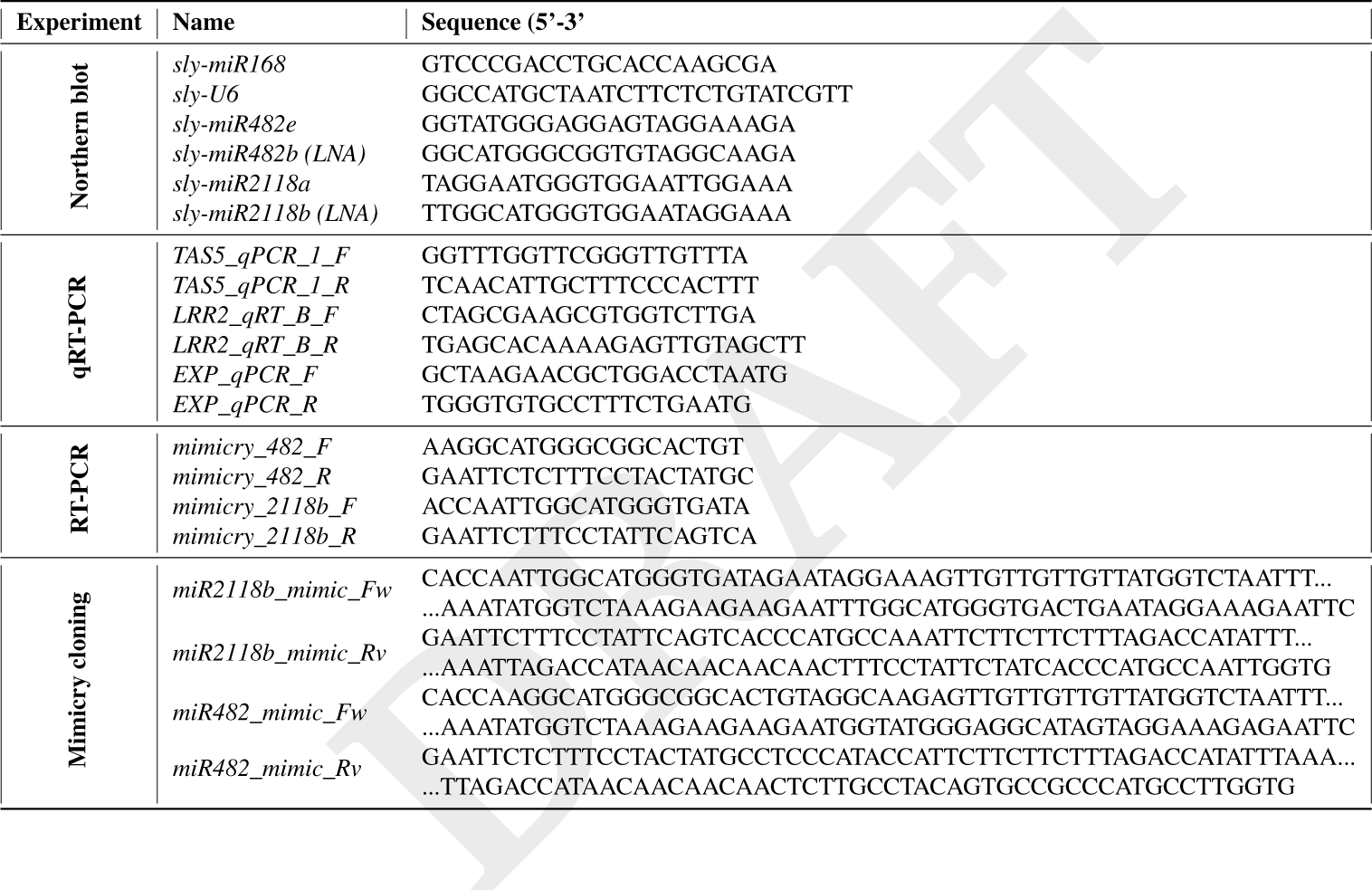
Oligonucleotide sequences used in this study.

